# Non-equilibrium cyclic conformations of DNA under electrokinetic confinement

**DOI:** 10.64898/2026.11.03.764104

**Authors:** Matheus A. S. Pessôa, Walter Reisner

## Abstract

Classic nanofluidic devices harness geometric confinement to modulate single polymer conformation and control chain dynamics. While classic nanofluidic devices have given rise to a range of applications, they are fundamentally limited by the static nature of their construction, which restricts real time control over molecular conformations. The reversible electrokinetic confinement (RECON) technique enables time-dependent control of DNA conformations in nanowells via dynamic modulation of field-induced confinement. Here, we demonstrate that applying dynamically varying confinement using stochastic and periodic signals can drive the confined polymer to access a range of non-equilibrium states, with the distinct configurations quantified via their radius of gyration tensor. For periodic driving, the device allows controlled cycling across conformational states, with distinct hysteresis observed in the shape anisotropy modulation. We quantify the interplay between the driving frequency, driving amplitude, and resulting hysteresis. Overall, our results establish RECON as a platform for active manipulation of single molecules, opening new avenues for studying non-equilibrium polymer physics and developing high-throughput, dynamically tunable nanofluidic devices.

## 1 Introduction

In a classic nanofluidic device, polymers such as dsDNA are squeezed into structures with dimensions smaller than the polymer’s equilibrium coil size in solution (characterized by the radius of gyration *R*_*g*_). In these confined environments, the interplay between conformational entropy [1, 2], excluded-volume interactions [3], and electrostatics [4–6] defines an effective free-energy landscape for the molecule, forcing the polymer to adopt equilibrium conformations determined by the geometry and spatial constraints of the confining structure. In nanoslits, where the molecule is confined between two parallel surfaces, confinement acts primarily in the direction normal to the slit plane, allowing the molecule to explore the two-dimensional plane of the slit. DNA consequently adopts flattened conformations whose in-plane size, anisotropy, diffusivity, and relaxation dynamics depend on the slit height, molecular contour length, and ionic conditions [7–10]. In nanochannels, the molecule is restricted in two transverse directions but remains free along the channel axis, suppressing transverse fluctuations and leading to DNA extension that forms the basis of a high-throughput genomic mapping approach [11]. Nanocavities, either fabricated individually or in arrays as embedded structures in a nanoslit, confine molecules completely along two lateral dimensions and partially along a third vertical dimension. Nanocavities can produce compact conformations or conformations partitioned between two or more cavities, with the exact induced conformation determined by the relationship between the cavity dimensions [12–15], nanocavity separation [12] and the equilibrium polymer size.

In classic nanofluidic devices, confinement is etched-in and static so that confinement control alone cannot be used to perform dynamic polymer manipulations. Externally imposed forces, implemented either hydrodynamically, electrically, or via tweezer-controlled sliding bead-gaskets, are required to perform the non-equilibrium manipulations required for polymer physics exploration and device operation. This includes forming folded [16], knotted [17], dynamically compressed polymer states [18–20] and transferring molecules from regions of low to high confinement [21]. As a step towards making nanofluidic devices more intrinsically dynamic, techniques have been developed to directly modulate device confinement by mechanical deflection of the device bonding lid, either via a piezo-controlled lens-pusher or pneumatic deflection [22–25]. While these approaches allow for more gentle loading of molecules into confined structures, and provide a higher degree of control over confinement, they require mechanical components that increase the timescales for confinement modulation. In addition, mechanical deflection tends to produce large-scale hydrodynamic flows arising from the mechanically displaced fluid. This prevents localized real-time modulation of the confinement landscape for exploring non-equilibrium effects that arise when confinement is modulated on timescales comparable to the polymer relaxation time.

Locally modulated electric forcing potentially provides a more versatile and dynamic means of inducing nonequilibrium conformations. The amplitude, direction, and temporal profile of electrically induced forces can be varied rapidly without the need to mechanically adjust the confining geometry. Electrical forcing can be used to induce molecular stretching [1, 26], collapse [27], folding [2, 16, 28], compression [29], and self-entanglement [30]. Tang et al. demonstrated that a uniform direct electric field can compress individual DNA molecules into compact, self-entangled conformations [30]. Electric fields have also been used to stretch knotted DNA [31] and regulate knot mobility through the tension imposed along the molecule [32], knot untying [33], and elucidate the mechanism of coil–stretch transitions [34]. Previous electrically driven approaches have primarily focused on pronounced conformational changes, demonstrating that electrical forcing can generate and regulate a broad range of nonequilibrium polymer conformations and dynamics. However, when the applied forcing varies in time, the molecular response is governed not only by the magnitude of the force but also by the competition between the driving timescale and the intrinsic relaxation times of the polymer.

The competition between driving timescale and polymer relaxation has been explored experimentally for single DNA chains using microfluidic systems to impose oscillatory extensional flow [35, 36]. The problem was recently examined theoretically and numerically for a single polymer subjected to an oscillatory pulling force [37]. When the driving period is long compared with the the polymer relaxation time, the chain follows the applied force approximately adiabatically and remains close to its instantaneous equilibrium conformation. As the forcing timescale approaches the the polymer relaxation time, the conformational response develops a phase lag relative to the applied force, producing hysteresis and energy dissipation over each driving cycle. At still higher frequencies, the polymer can no longer reorganize globally within a single cycle, and only faster, short-wavelength conformational modes can respond appreciably.

Dynamically tunable the confinement landscapes may provide both simplified models for fluctuating biophysical environments and a platform for programmable single-molecule manipulation. Building on the picture of dynamically driven polymer relaxation, an open question is whether comparable non-equilibrium conformational responses can be generated and continuously regulated by modulating the the confinement landscape itself, rather than by changing the physical geometry of the device or applying an external flow. Recently, we demonstrated that sculpting electric fields in nanocavities using high frequency, low-voltage fixed-polarity signals leads to high-throughput molecular confinement, release, and real time manipulation of confined nanoentities such as dsDNA, liposomes, and nanotubes [38]. Here, we confine DNA molecules in nanowells using the electrokinetic confinement device and apply real time confinement modulation to study molecular shape dynamics; we demonstrate that electrokinetic confinement enables control of non-equilibrium conformations in real time following user-defined dynamics. Con-formational shape dynamics are accessed by measuring the gyration tensor **G**(*t*) of the confined molecules. Through **G**(*t*), we obtain dynamical information about the in-plane projection ellipse of the confined molecule, with major (*R*_*M*_) and minor (*R*_*m*_) axes. We then probe the influence of different electrical landscapes on conformational changes in real time, visualizing the molecular dynamics using trajectory plots in *R*_*M*_ − *R*_*m*_ conformational phase space. First, we explore the simplest confinement scenario in which the confinement waveform has a fixed voltage and study the influence of waveform frequency and nanowell geometry (by varying the radius of the nanowells). We then explore more complex waveforms, using stochastic driving and amplitude modulation of the confinement waveform. We find that stochastic driving leads to greater excursions from the nanowells. Driving the system using periodic amplitude modulation leads to cyclic trajectories in the phase space, which we probe by varying the amplitude-modulation depth and modulation frequency. Measuring the trajectories in the conformational phase space, we observe hysteresis in the polymer conformational dynamics during the driving cycles. To obtain analytical predictions for the shape of the conformational cycles and rationalize the origin of hysteresis, we propose a coarse-grained free-energy landscape model based on the confinement of a DNA molecule within a cylindrical cavity under the influence of an external sinusoidal driving force. We find that the cyclic trajectories in the conformational phase space are related to the differing relaxation timescales of the major and minor axes, *R*_*m*_ and *R*_*M*_.

## 2 Methods

### 2.1 Experiments

The device is based on a parallel-plate geometry. The top electrode is an indium tin oxide (ITO)-coated coverslip (resistivity *ρ* = 15−30 Ω, and thickness 0.13−0.17 mm). The bottom electrode contains an array of nanowells with embedded electrodes, following the same workflow reported in [38]. Starting with an ITO-coated wafer, we deposit a 400 nm-thick layer of silicon nitride via plasma-enhanced chemical vapour deposition (PECVD). The nanowells are then defined via electron-beam lithography and etched through the nitride using reactive ion etching (RIE). At the end of the nanofabrication process, a 2 nm passivating layer of titanium dioxide (TiO_2_) is deposited on the surface of the chips via atomic layer deposition (ALD). The top electrode is separated from the cavities by a 30 *µ*m patterned microfluidic tape that defines the flow cell.

To initiate an experiment, we introduce 1× tris/boric acid/ethylenediaminetetraacetic acid (EDTA) buffer solution with *λ*-DNA into the flow cell. The molecules are fluorescently labeled with YOYO-1 using a 1:10 dye-to-base-pair ratio and pre-diluted 10–100 times prior to the experiment (the DNA contour *L* = 21 *µ*m for YOYO-1-labeled *λ*-DNA at this dye ratio). We also add 2% *β*-mercaptoethanol (BME) to the buffer to suppress DNA photobleaching and photodamage [39]. The chip is connected externally to a wave-form generator via a silver wire, soldered to copper electrical contacts with jumper wires, which are then connected to the generator. For imaging, we use a 100× widefield oil-immersion objective (Nikon CFI Apo 100×, N.A. 1.4) and a Teledyne Photometrics 95B camera (95% efficiency). We use a home-built setup for laser excitation (*λ* = 525 nm) and a Texas Red FITC filter set. Approximately 3–4 *µ*L of diluted DNA solution is introduced into the flow cell through a manually drilled inlet on the chip backside.

The top and bottom plates are connected in series with a waveform generator. The bottom electrode where the etched nanowells are defined is biased positively (Fig.1A), while the top plate is grounded. Applying an offset voltage between the two plates induces an alternating current with fixed polarity (ACfp) through the bottom electrode, leading to confinement of molecules in the nanowells. The magnitude of the applied current is ~ 4pA*/µ*m^2^ at 3 V, as previously measured in [38]. Upon application of the ACfp, an electric field funnel is formed within the cavities, leading to capture of negatively charged DNA in the nanowells and confinement within the boundaries of the cylindrical volume. Once the voltage is zeroed, the molecules are released in solution and diffuse freely in bulk (Figure 1B).

**Figure 1:**
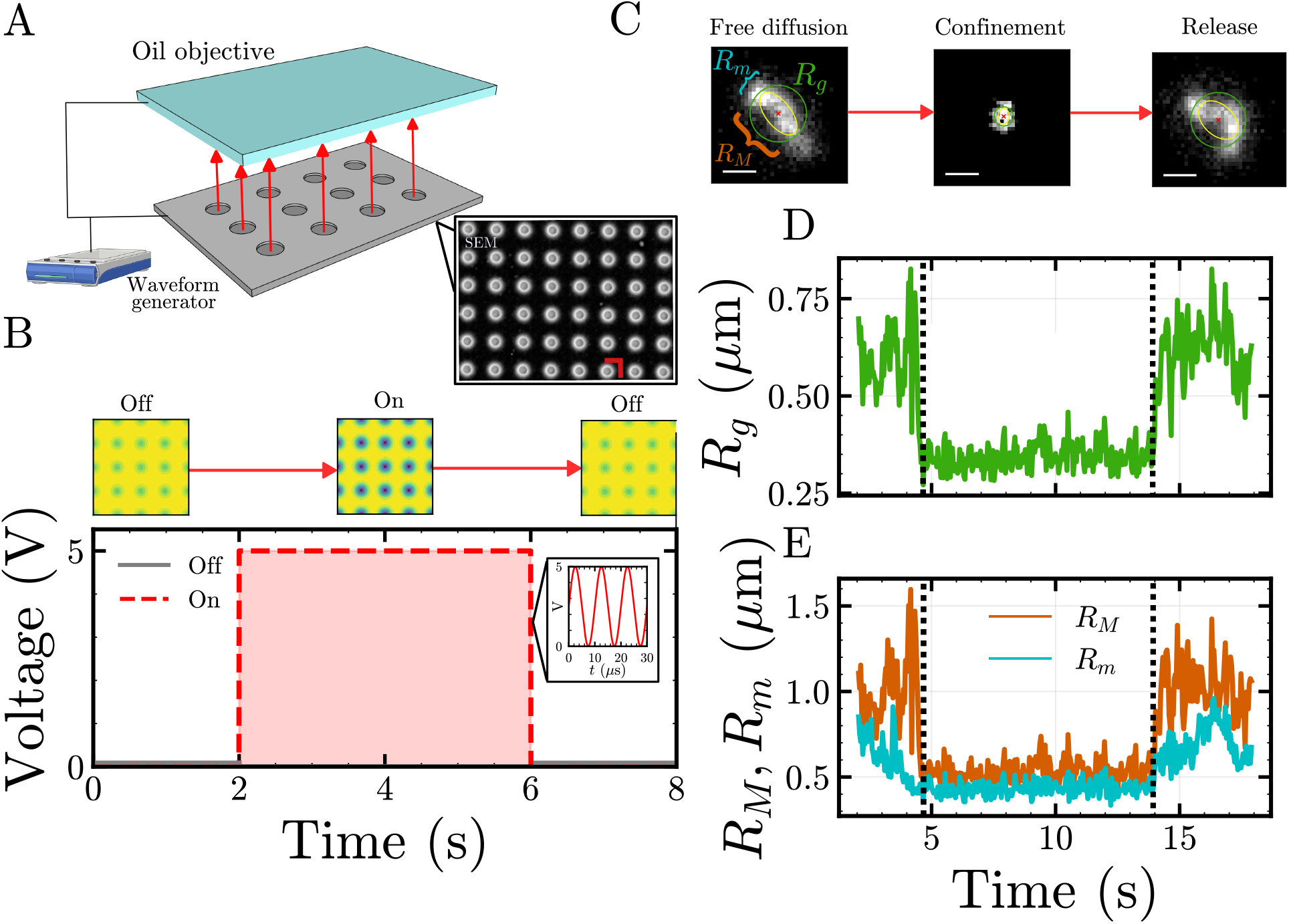
A) The reversible electrokinetic device is based on a parallel-plate configuration, with a top plate consisting of a uniform indium tin oxide (ITO) electrode, and a bottom plate consisting of etched ITO electrode wells, connected externally to a waveform generator. The nanocavities are etched through a silicon nitride layer, and are uniformly spaced (2 *µ*m), as shown in the SEM. The imaging is performed through a top transparent ITO electrode, using an inverted microscope. The arrows give the direction of the electric field. (B) A synthetic potential landscape for molecule capture is toggled on when a fixed polarity AC signal (ACfp) is input in the system; the landscape toggled off when the ACfp signal is zeroed. (C) *λ*-DNA initially freely diffuses in bulk solution, then is confined in the nanowell upon application of the ACfp signal, and is released when the signal is zeroed. The molecular radius of gyration is shown as a yellow ellipse, and the gyration-tensor axes in blue. The scale bars correspond to 1 *µ*m. (D) Plot of gyration radius *R*_*g*_ and (E) both the major (*R*_*M*_) and minor (*R*_*m*_) axis of the gyration tensor as a function of time. When field confinement is initiated, the molecular dimensions are reduced as the molecule is squeezed to fit within the nanowells.

Upon application of a high frequency ACfp sinusoidal voltage signal to all wells in the array, we enable capture of tens to hundreds of molecules in the ≈ 100 *µ*m-wide microscope field of view and their confinement until the signal is zeroed. The applied potential *ϕ*(*t*) can be described as

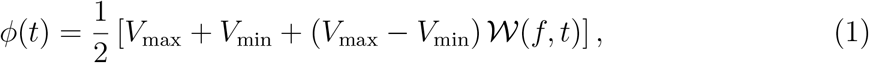

with a minimum voltage *V*_min_, maximum voltage *V*_max_, and frequency-dependent waveform *W*(*f, t*). In Fig. 1, for example, *V*_min_ = 0 V, *V*_max_ = 5 V and *f* = 100 kHz. Unless otherwise stated, we will utilize a sine wave with offset as described in Eq. 1 (i.e. W(*f, t*) = sin(2*πft*)). The electrical confinement landscape therefore can be defined by the user by varying the applied waveform, voltage frequency, and voltage bias.

### 2.2 Image analysis

The DNA fluorescence micrographs give access to the two-dimensional projection of the total conformation, including information on shape anisotropy and overall coil size. First, areas of interest containing the confined molecules are selected in the acquired image stacks, with area *N* = *i* × *j* pixels. The regions of interest are cropped from the main stack, saved, and noise is subtracted using the algorithm described in [40]. We compute the center-of-mass (CoM) of the molecule at each frame using

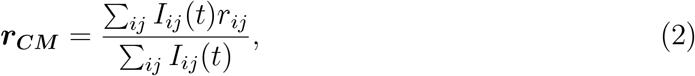

where *I*(*r, t*) = *I*_*ij*_(*t*) is the intensity of the pixel (*i, j*) at a given time *t*, and *r*_*ij*_ denotes the pixel center. We can access anisotropy in the DNA shape, i.e., differences in the coil extension along axes parallel and perpendicular to the axis of longest extension, by computing the covariance matrix of the intensity distribution, or gyration tensor

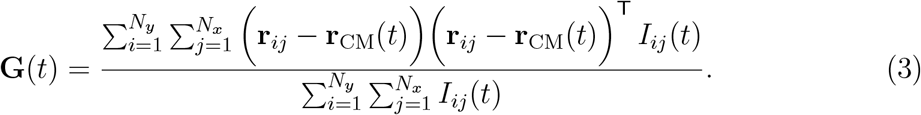

The gyration tensor defines an ellipse with major radius *R*_*M*_ and minor radius *R*_*m*_. The radii are obtained by taking the eigenvalues of the gyration tensor (Figure 1 C),

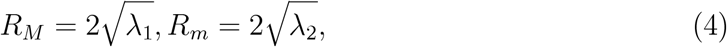

where *λ*_1_ and *λ*_2_ are the principal components of **G**. Figure 1C shows the principal radii of an example molecule during free diffusion, confinement, and release. We can also compute the radius of gyration as

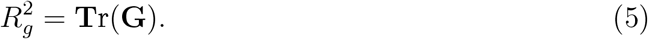

We further define the projection of *R*_*g*_ in 2-D, parallel to the confining plane, as

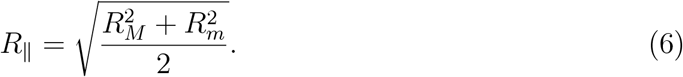

For each experiment we compute the gyration tensor and extract *R*_*M*_ and *R*_*m*_ on a frame-by-frame basis, giving access to the coil shape dynamics as a function of time (Figure 1D,E). This approach for bulk DNA measured in free solution leads to an *R*_*g*_ ≈ 0.78 ± 0.15 *µ*m, comparable to the accepted literature value obtained from diffusion (0.73 *µ*m [41]). Note that equilibrium polymer coils have a well-known intrinsic shape anisotropy leading to distinct principal axes even in bulk [42], so we expect that in general *R*_*M*_ *> R*_*m*_. To quantify the timescales associated with anisotropy changes, we can take the auto-correlation function of 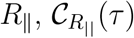, as

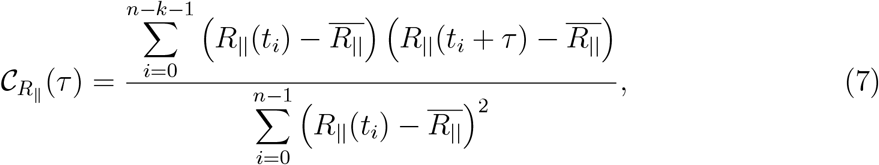

with *τ* = *k*Δ*t*, where Δ*t* is the frame duration, in seconds, and *k* is the integer number of frames. In this equation, *R*_*M*_ (*t*) represents the value of the major axis at each time point, and 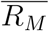 represents its average value throughout the time series. We use the same form of the autocorrelation function described in Eq. 7 to analyze the experimental results.

## 3 Results

### 3.1 Confinement anisotropy under constant amplitude driving for variable well radii

First, we investigate anisotropy in polymer shape for conditions of constant-amplitude sinusoidal driving (Eq. 1) using wells with variable radii (*R*_well_ = 250, 300, 350, and 400 nm). We quantify the coil size and shape anisotropy across the fluorescence recording via the approach outlined in the previous section. The gyration tensor ***G*** (Eq. 3) computed for each frame defines the projected two-dimensional ellipse characterizing the polymer coil for each time point. The major and minor axes (Eq. 4) of the ellipse together characterize the coil shape anisotropy; the radius of gyration *R*_*∥*_ (Eq. 6) of the projected ellipse characterizes the coil size.

Upon confinement, the molecule is compressed due to the larger size of the free solution DNA coil relative to the dimension of the confining nanowells (*R*_well_ *< R*_*g*_). Figure 2A shows the distributions of the major and minor axes, *R*_*M*_ and *R*_*m*_, for different well radii (*f* = 100 kHz, *V*_min_ = 0 V, *V*_max_ = 5.0 V). Smaller wells produce a wider distribution of molecular conformations with greater anisotropy as less of the molecule is confined entirely within the well. Greater anisotropy is expected when less of the molecule is confined because the unconfined portion of the DNA adopts the more anisotropic configurations expected for bulk coils. Increasing the well radius progressively suppresses these fluctuations, resulting in more compact isotropic conformations that have a more similar distributions of *R*_*m*_ and *R*_*M*_.

**Figure 2:**
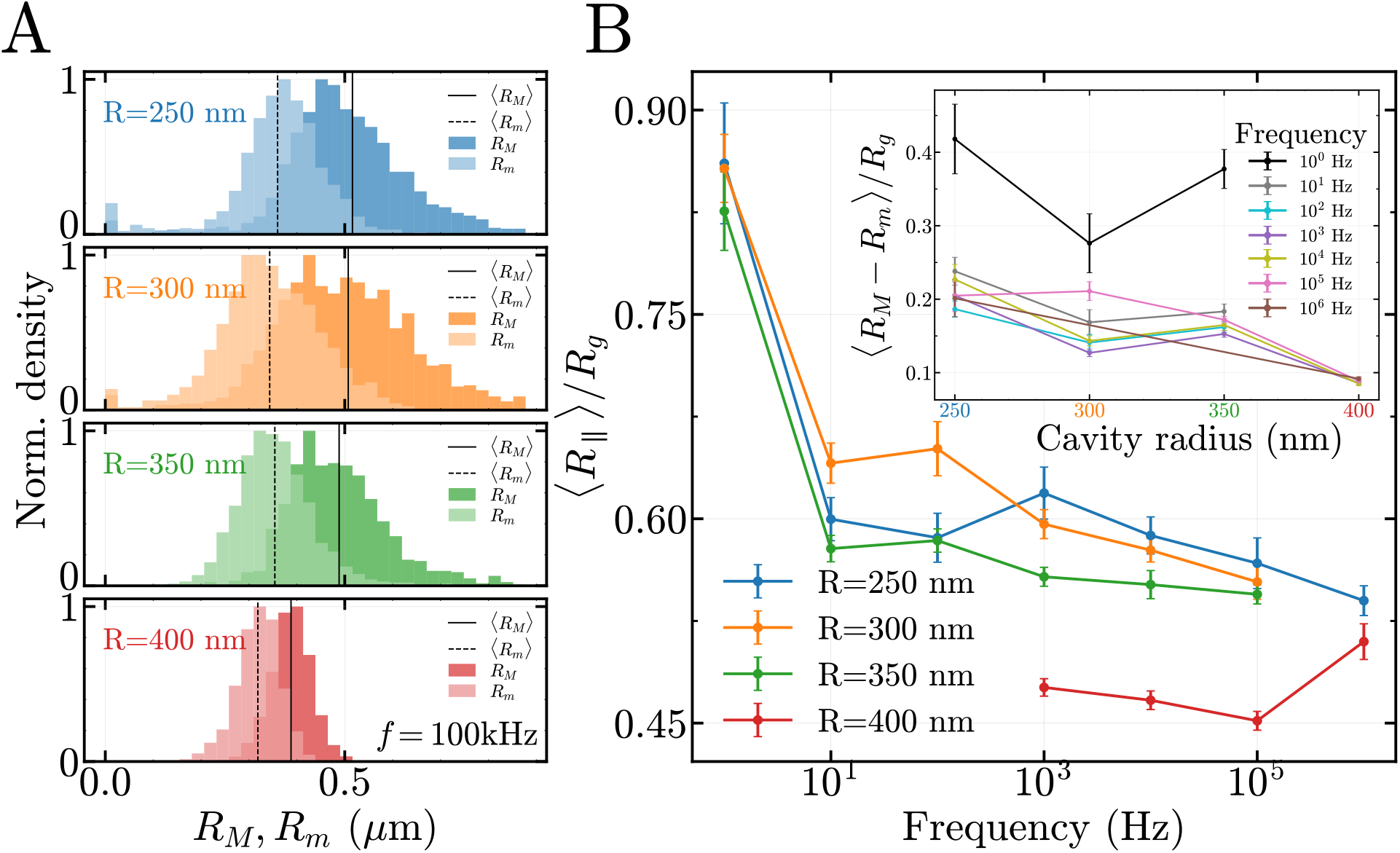
A) Histograms showing the major and minor axes of the radius of gyration tensor, for different well radii and fixed frequency and voltage (*f* = 100 kHz, *V*_min_ = 0 V, *V*_max_ = 5.0 V), with the dashed and outlined vertical lines giving the averaged principal radii. B) The average values of *R*_*∥*_ as a function of frequency for different well radii. The value of *R*_*∥*_ is scaled by the value of *R*_*g*_ in bulk. The inset shows the difference between the averages of *R*_*M*_ and *R*_*m*_ as a function of well radius, at different frequencies, also normalized by *R*_*g*_.

We also investigate how frequency affects shape anisotropy and coil size (Figure 2C). For this purpose, we study the distribution of *R*_*M*_ and *R*_*m*_ (presented in Supplementary Material 8.1). At low frequency (1 Hz), the distributions are broader and the average value *R*_*∥*_ approaches the measured *R*_*g*_ in bulk. Pronounced changes in the distribution of *R*_*∥*_ occur in the transition from low (1 Hz) to intermediate (*>* 10 Hz) frequencies. This can be understood by comparing the period of the driving signal, *T* = 1*/f*, to the time for the molecule to diffuse across the vertical dimension of the well. The diffusion time can be estimated as *τ*_*d*_ *≃ h*^2^*/D*_DNA_ ≈ 0.18s, with *D*_DNA_ = 0.5 *µ*m^2^*/*s, and a cavity height *h* = 400 nm. When the driving period satisfies *T > τ*_*d*_, the molecule can diffuse out of the nanowell vicinity during an oscillation cycle, leading to a less compressed configuration where *R*_*m*_ and *R*_*M*_ approach their values in the absence of confinement. At frequencies above 10 Hz, the distribution of *R*_*∥*_ becomes tighter, with differences comparable to the standard deviation of the mean.

### 3.2 Influence of Waveform

The RECON approach enables investigation of how different waveforms, such as periodic ACfp, stochastic and amplitude-modulated ACfp, influence the confinement dynamics (Fig. 3). The effect of these waveforms can be conveniently viewed in the (*R*_*M*_, *R*_*m*_) plane, defining a coarse-grained conformational space (Figure 3A). In this space, anisotropy leads to a displacement of the configuration beneath the *R*_*m*_ = *R*_*M*_ unity line. A constant-amplitude sinusoidal ACfp is quite tightly localized around a central average (Fig. 3 A, left panel, gives compiled results for *n* = 12 molecules). The autocorrelation function C_*R∥*_(*τ*) for the constant-amplitude sinusoidal ACfp exhibits no characteristic oscillatory or time-dependent signature (Fig. 3B,C). Note that the relaxation of spontaneous conformational fluctuations in this case occurs at a timescale too small to resolve given the time step used (0.042 s). Similar behavior is observed for the other well sizes investigated, as shown in Supplementary Material 8.2. Essentially, once confined using a fixed sinusoidal waveform at high frequency, the DNA molecules acquire largely static conformations subject to small fluctuations about the mean.

**Figure 3:**
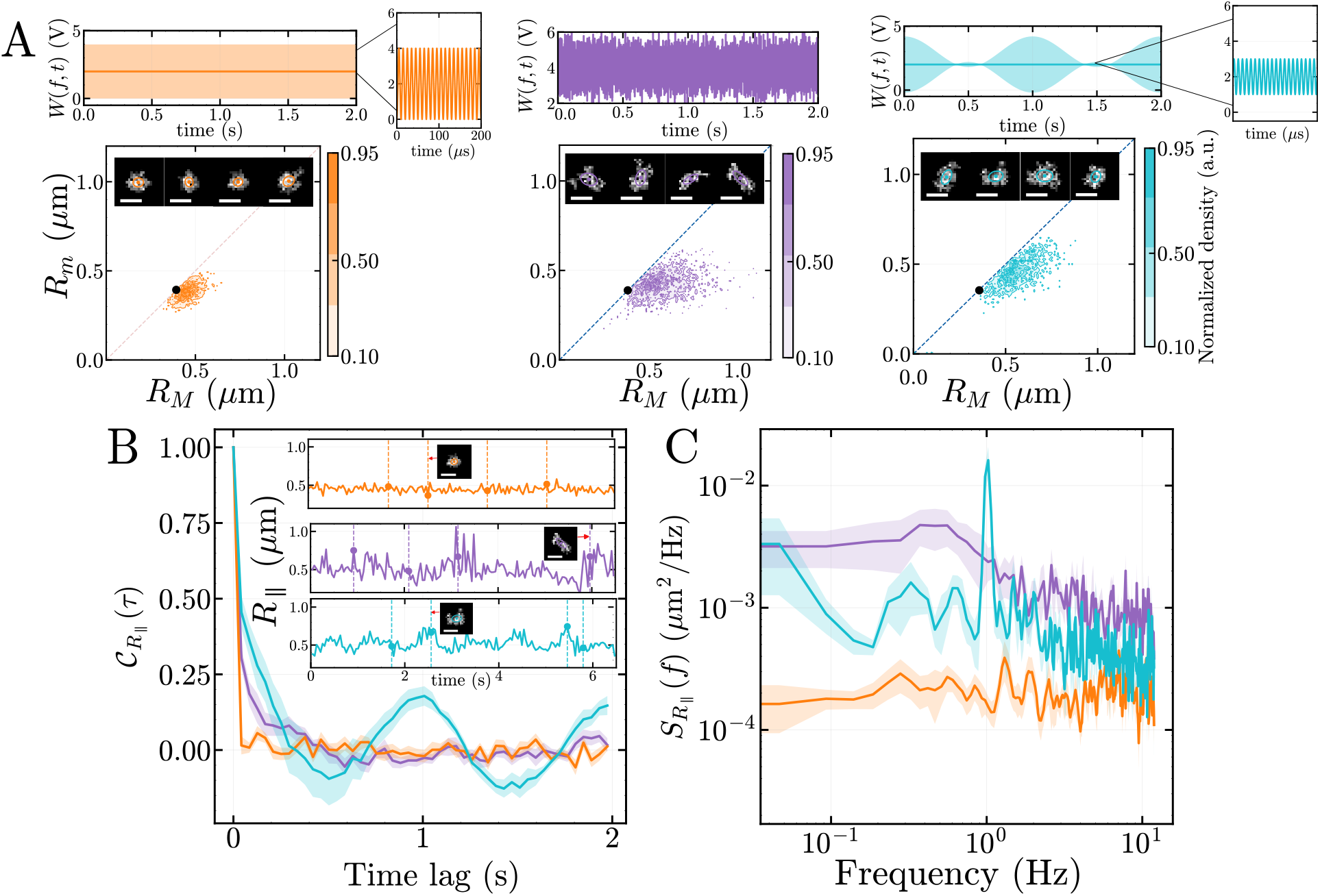
A) Comparison of constant amplitude sinusoidal signal (orange, left, *n* = 12), stochastic (purple, center, *n* = 9), and sinusoidal signal with sinusoidal amplitude modulation (cyan, right, *n* = 6) at fixed cavity geometry (*r*_cavity_ = 400 nm). The upper row shows the applied signals, while the lower row shows the corresponding conformational distributions in the (*R*_*M*_, *R*_*m*_) plane, where *R*_*M*_ and *R*_*m*_ are the major and minor conformational radii, respectively. The black dots are positioned at the radius of the cavity, *R* = 0.4 *µm*, and the insets show representative molecular conformations. A constant sinusoidal waveform gives rise to a more localized distribution, the stochastic waveform broadens the range of accessible conformations, and a sinusoidal waveform with sinusoidal amplitude modulation produces a structured distribution associated with periodic conformational cycling. (B) Autocorrelation function 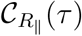 for the three waveforms. The insets show representative *R*_*∥*_(*t*) trajectories and corresponding molecular images. The relaxation time for the stochastic signal is *τ*_*R*_ = 0.15 ± 0.01s, obtained via exponential fit. Constant amplitude sinusoidal and stochastic waveforms produce predominantly nonperiodic fluctuations, whereas sinusoidal amplitude modulation generates a clear oscillatory correlation at the imposed modulation period. Shaded regions indicate the variability between measurements. (C) Power spectral density 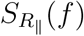 of the corresponding conformational trajectories. Sinusoidal modulation produces a pronounced spectral peak at the driving frequency, whereas the constant amplitude sinusoid yields a broadband spectrum without a comparable deterministic frequency signature. The stochastic waveform induces conformational relaxation at a larger timescale (~ 1 s) that can be resolved given the experimental time step.

In contrast, time-dependent stochastic waveforms provide a means of continuously perturbing the effective free-energy landscape experienced by the confined molecule. RECON can induce randomly fluctuating free-energy landscapes through the application of stochastic voltage signals [38]. Noise-driven modulation of DNA dynamics has been explored in related nanofluidic systems, such as electrokinetically assisted barrier-crossing events induced via electrokinetic noise [43]. This work shows shows that increasing the variance of the applied electrokinetic noise broadens the accessible range of molecular motion and enhances DNA diffusion [44], behavior that can be interpreted in terms of a noise-dependent effective temperature. In the middle panel of Figure 3A, we show that stochastic voltage modulation similarly broadens the conformational distribution of DNA confined within a well (results shown for *n* = 9 molecules). Our signal utilizes a Gaussian-distributed waveform with an offset so the signal fluctuates randomly around a prescribed mean voltage *V*_0_. Note that the offset was sufficiently high to ensure that the molecule is confined throughout the experiment. Lowering the offset can lead to polymer hopping or transitioning between different cavities [38].

Under stochastic driving, we observe irregular excursions of the polymer within the well, which lead to more elongated molecular conformations and appear to lead to stronger correlations in *R*_*∥*_. In this case, 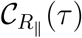 decays exponentially without a persistent recurring signature (Fig. 3B), consistent with the stochastic character of the applied forcing, and the corresponding power spectral density remains distributed over a broad range of frequencies with a clear relaxation corner frequency (Fig. 3C). The effective correlation timescale obtained from fitting an exponential relaxation is *τ*_noise_ ~ 0.15 ± 0.01 s, sufficiently high to be resolved given the time step.

Although molecular confinement can be achieved using a fixed-amplitude signal at low frequencies, the lower frequency limit is constrained by the prolonged duration of each voltage half-cycle (e.g. for a 1 Hz cycle, the positive bias lasts for 500 ms). This can promote dielectric breakdown and electrolysis leading to bubble formation in the device. An alternative approach to achieve molecular confinement while recovering the low-frequency dynamics is to dynamically modulate the voltage waveform [38]. Using the general wave-form representation introduced in Equation 1, we define the amplitude-modulated (AM) waveform as *W*_AM_(*t*) = sin(2*πf*_*m*_*t*) sin(2*πf*_*c*_*t*) where *f*_*m*_ is the modulation frequency defining the frequency of oscillation of the confinement free energy landscape and *f*_*c*_ is the carrier frequency. Low-frequency AM allows cycling between deeper and shallower potential wells form molecular confinement. On the right panel of Figure 3A, we show the conformational distribution under amplitude modulation at *f*_*m*_ = 1 Hz (results shown for *n* = 7 molecules). Compared to stochastic forcing (Fig. 3A, middle panel), sinusoidal amplitude modulation produces a more defined pathway in conformational space (Fig. 3A, right panel), reflecting the reproducible cycling of the molecule between states associated with weaker and stronger field-induced funnel forces. This is apparent as clear sinusoidal modulation in 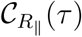 (Fig. 3B) and a prominent peak in the power spectral density at the driving frequency of 1 Hz (Fig. 3C).

### 3.3 Conformational cycling via periodic modulation of potential well depth

#### 3.3.1 Effect of varying modulation depth

Amplitude modulation (AM) is a general approach for exerting in situ control over molecular conformation in real time. Figure 4 shows how modulating the voltage between a maximum (I) and a minimum (II) leads to molecular excursions from the trap, which we quantify via measurements of *R*_*M*_, *R*_*m*_ and *R*_*∥*_. The degree of amplitude modulation can be quantified by introducing the percentage AM modulation depth:

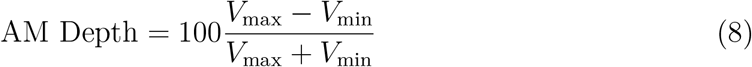

**Figure 4:**
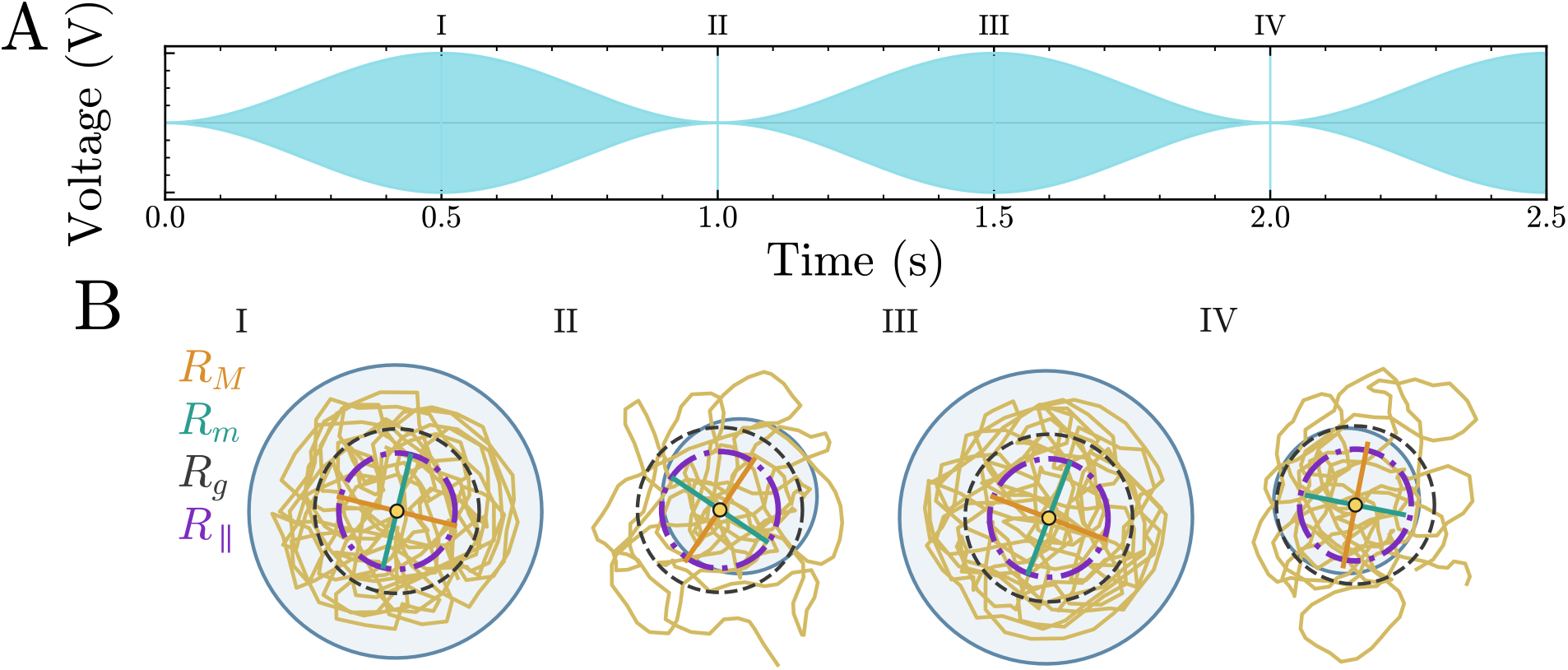
A) Amplitude-modulated voltage and B) cartoon depictions of representative conformational states of the confined DNA. When the voltage is maximal (I, III), the *R*_*∥*_ is minimal, and when the voltage is minimal, *R*_*∥*_ is maximized.

This quantifies the extent of the waveform that is dynamically modulated (Fig. 12A).

To quantify the molecular dynamics under AM modulation, we determine the averaged dynamic trajectory of a molecule over a cycle in *R*_*M*_ –*R*_*m*_ conformational phase space (Fig. 5). To achieve this, we extract *R*_*∥*_ for each molecule at a given modulation depth and temporally align the individual traces such that equivalent phases of the modulation cycle coincide. The aligned trajectories are then divided into complete modulation periods, allowing each cycle of each molecule to be treated as an independent realization of the conformational response, which can then be averaged. The highlighted points MIN 1, MAX, and MIN 2 in Fig. 5A identify the characteristic conformational states reached during the cycle. MIN 1 corresponds to the initially compact configuration at *V*_max_ under the strongest confinement, MAX marks the state of largest conformational extension as the confinement weakens at *V*_min_, and MIN 2 corresponds to the subsequent return to a more compact configuration as the confinement increases. Figure 5B shows the phase-space trajectory arising from the average of *n* = 10 cycles. We observe that hysteresis is present in the cycle, with the averaged trajectory being traversed clockwise with distinct forward (MIN-1 to MAX) and backward (MAX to MIN-2) portions; this corresponds to *R*_*m*_ leading and *R*_*M*_ lagging (Figure 5C).

**Figure 5:**
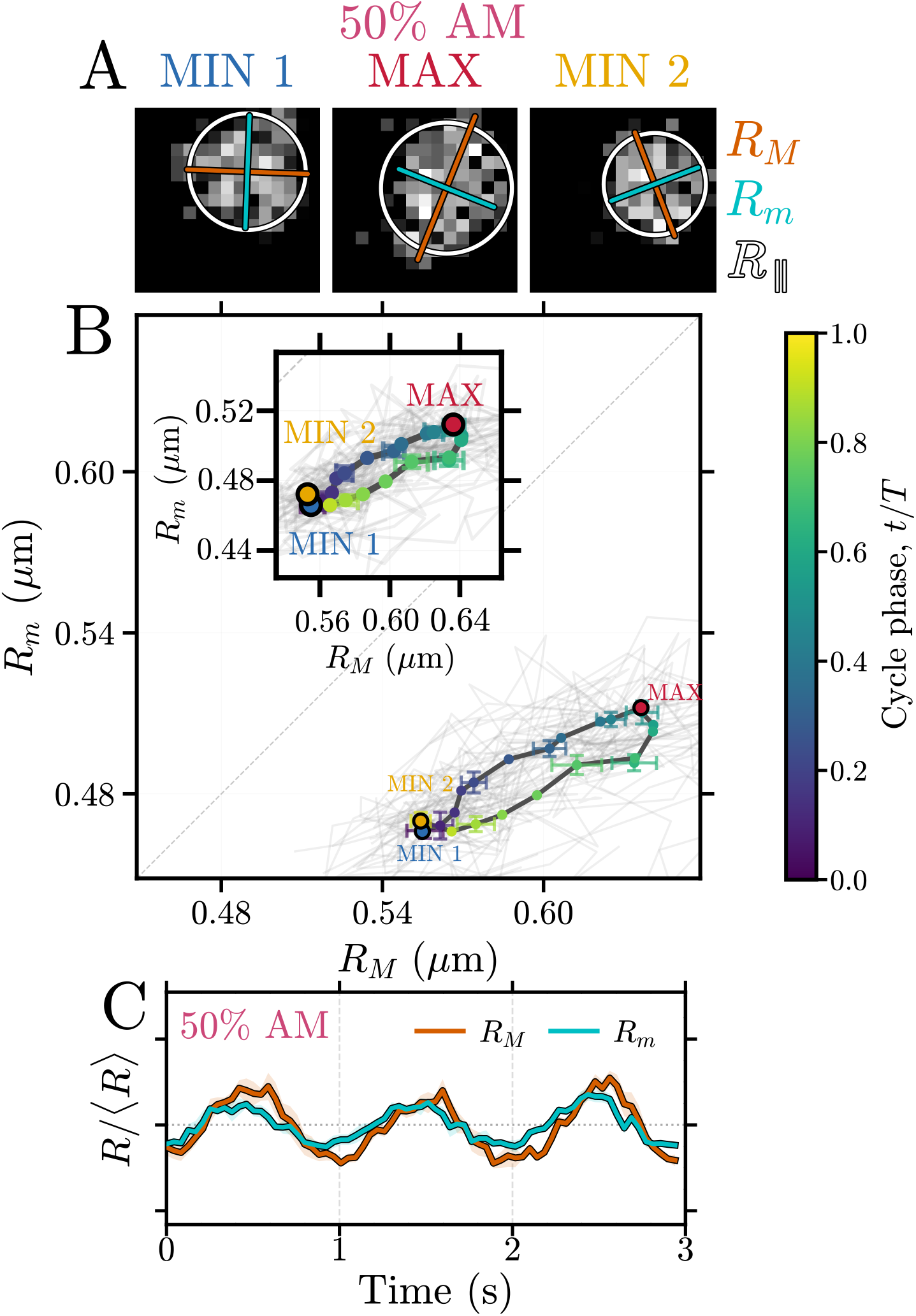
Conformational response of DNA confined in an electrokinetic well under amplitude-modulated electrokinetic confinement for 50% modulation depth. A) Representative fluorescence images at the starting point in the cycle where the signal is maximized and the coil size is minimized (MIN 1), the cycle mid-point where the signal is minimized and coil size is at its maximum (MAX), and the return to a state of minimum coil size (MIN 2). B) Corresponding conformational trajectories in *R*_*M*_ –*R*_*m*_ phase space for *n* = 10 cycles along with the averaged trajectory (pale gray indicates underlying single-cycle trajectories; black bold curve indicates average). Colored points along the averaged trajectory indicate progression through the modulation cycle according to the normalized cycle phase *t/T*, with dark blue indicating the start of the cycle, red the middle maximum point on the cycle and bright yellow indicating the cycle end. Error bars correspond to standard error of the mean of each point within the cycle and indicate the variability of the ensemble-averaged conformational coordinates at these positions. Inset shows a magnified view of the phase-space trajectory. C) Time series of *R*_*M*_ –*R*_*m*_ dynamics with both quantities normalized to their average values. Note that *R*_*m*_ leads and *R*_*M*_ lags.

The AM depth was then systematically varied, with the carrier frequency kept fixed at *f*_*c*_ = 100 kHz and modulation frequency held at *f*_*m*_ = 1 Hz (Figure 12A). For easier comparison, Figure 12B shows the phase-space trajectories in normalized coordinates (*R*_*M*_ */⟨R*_*M*_ *⟩* and *R*_*m*_*/⟨R*_*m*_*⟩*). At low modulation depths, between 10% and 30%, the molecules undergo limited excursions in the *R*_*M*_ –*R*_*m*_ conformational phase space and remain close to their mean confined conformation throughout the cycle. This suggests that the molecule is primarily confined in the well and does not undergo strong periodic excursions. As the modulation depth is increased to 50%, cyclic trajectories exhibiting hysteresis become apparent.

We also explore the effect of varying modulation frequency. Figure 7 shows the same analysis applied for modulation frequencies ranging from 1–5 Hz using a fixed modulation depth of 120%. The degree of hysteresis and overall cycle excursion diminish as the frequency is increased. This effect can be quantified by determining the area of the phase-space loop by fitting the experimentally determined averaged trajectory to an elliptical path.

## 4 Origin of Hysteresis in Conformational Cycling

We propose that the observed hysteresis in conformational cycling arises from the different relaxation timescales associated with the major and minor axes of the polymer coil. When the coil is almost entirely confined in the well at constant amplitude driving (e.g. Fig. 3A, left panel) or nearly constant amplitude driving (Fig. 6, 10% Modulation Depth), then *R*_*m*_ and *R*_*M*_ are centered at and tightly distributed around the well radius of 400 nm. When the modulation depth increases, polymer contour can escape the well during the downward voltage swings, entering the region of solution above the well where it will expand outward to follow approximately bulk polymer statistics. We expect hydrodynamic friction to differ for polymer expansion along the minor and major axes, with the friction highest for expansion along the major axis, leading to slower dynamics along the major axis. In addition, we expect entropic spring constants for deformation along the principal axes *R*_*i*_ to scale as 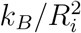, so that the major axis is softer, further increasing its relaxation time relative to the minor axis. This suggests that minor axis dynamics should lead, and major axis dynamics should lag, leading to the observed hysteresis.

**Figure 6:**
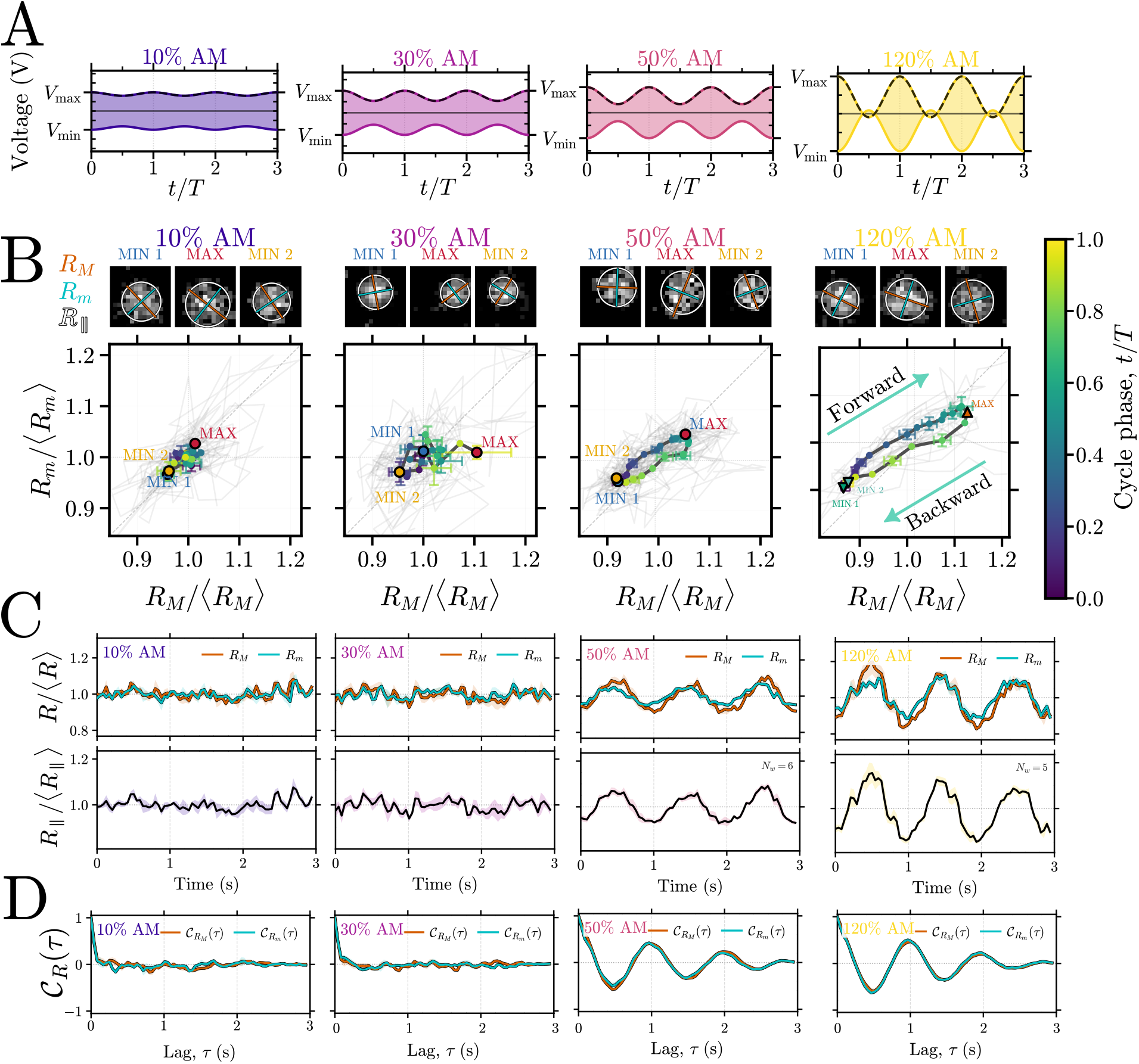
Amplitude-dependent conformational response of confined DNA under amplitude-modulated electrokinetic confinement. A Applied voltage waveforms for modulation depths of 10%, 20%, 30%, 50%, 100%, 110%, and 120%. The upper and lower envelopes indicate the instantaneous modulation limits, *V*_max_ and *V*_min_, over three modulation periods, *t/T*. B Corresponding conformational trajectories in the normalized *R*_*M*_ –*R*_*m*_ phase space for *n* = 10 cycles along with the averaged trajectory (pale gray indicates underlying single-cycle trajectories; black bold curve indicates average). Representative fluorescence images are shown at the starting point in the cycle (MIN 1), at the cycle mid-point (MAX) and at the return to a state of minimum coil size (MIN 2). Colored points along the averaged trajectory indicate progression through the modulation cycle according to the normalized cycle phase *t/T*, with dark blue indicating the start of the cycle, red the middle maximum point on the cycle and bright yellow indicating the cycle end. Error bars correspond to standard error of the mean of each point within the cycle and indicate the variability of the ensemble-averaged conformational coordinates at these positions. Increasing modulation depth produces progressively larger excursions in *R*_*M*_ */⟨R*_*M*_ *⟩* and *R*_*m*_*/⟨R*_*m*_*⟩*, with the forward and backward portions of the cycle defining a closed conformational trajectory. C) Time series plots of *R*_*M*_, *R*_*m*_ and *R*_*∥*_ and D) Autocorrelation function that clearly shows the transition to periodically modulated behavior at 50% modulation depth. For 10% modulation depth, the relaxation timescales are *τ*_*R*_ ~ 0.1 s, while for 30%, *τ*_*R*_ ~ 0.4 s

**Figure 7:**
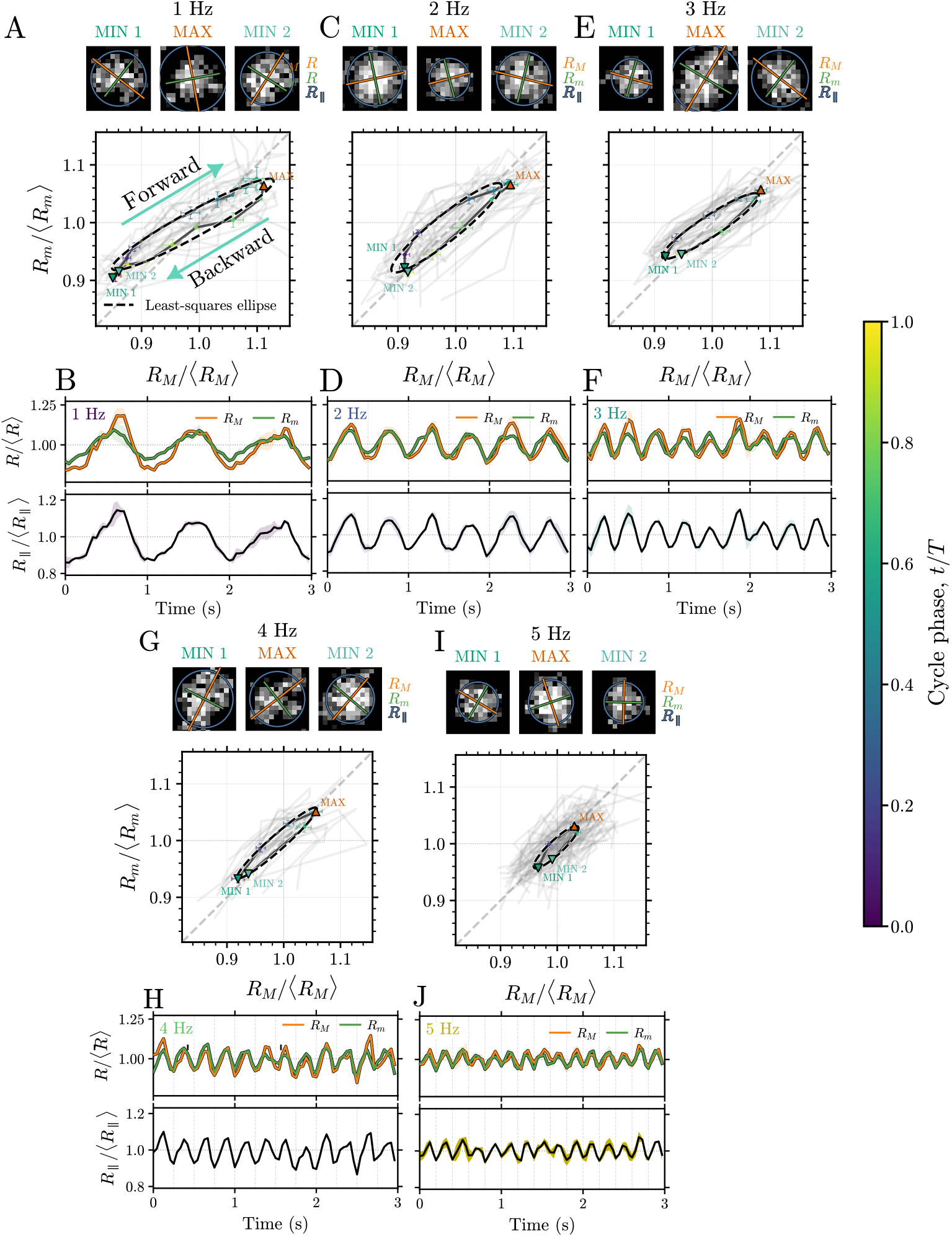
Conformational cycles of *R*_*M*_ */⟨R*_*M*_ *⟩* and *R*_*m*_*/⟨R*_*m*_*⟩* under amplitude modulation at 1–5 Hz. In panels A, C, E, G, and I, the trajectories are color-coded by the normalized cycle phase, *t/T*, evolving from the first minimum state (MIN 1), through the maximum state (MAX), and back to the second minimum state (MIN 2). Experimental fluorescence images are shown above the trajectories, with *R*_*M*_, *R*_*m*_, and *R*_*∥*_ overlaid. Individual experimental cycle trajectories are shown in light gray, while the ensemble-averaged trajectory is highlighted. The gray diagonal denotes *R*_*M*_ = *R*_*m*_, while the arrows in A indicate the forward and backward directions of evolution through the modulation cycle. Dashed black curves indicate the least-squares ellipse fits to the phase-space trajectories. Panels B, D, F, H, and J show the time traces over 3 s. The upper subpanels show *R*_*M*_ */⟨R*_*M*_ *⟩* and *R*_*m*_*/⟨R*_*m*_*⟩*, while the lower subpanels show *R*_*∥*_*/⟨R*_*∥*_*⟩*, with the shaded regions indicating the variability of the ensemble-averaged response. The trajectories become progressively shorter with increasing frequency, demonstrating the attenuation of the conformational excursion as the modulation period becomes shorter relative to the molecular reorganization time.

To quantify this picture, we develop a simplified semi-quantitative polymer model that considers the dynamics of contour exchange between inside and outside the well coupled to the expansion dynamics of the coil. To model the contour exchange dynamics, we adopt an approach developed originally for nanocavities embedded in a nanoslit [12, 14]. In this approach, we consider a polymer with contour *L*_in_ inside the well, contour *L*_out_ outside the well, and total contour *L* so that *L* = *L*_in_ + *L*_out_. Note that, without invalidating this approach, we can generalize the notion of contour confined in the well to include the contour confined in the well and confined in the portions of electric field funnel immediately adjacent to the well where the confining potential is strong (exceeds thermal energy).

The polymer confined in the well has a free energy *F*_well_*/k*_*B*_*T* arising from confinement-induced entropy loss, which scales linearly with *L*_in_ (*A*_conf_*L*_in_) and self-exclusion, which scales quadratically with 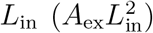. The prefactors depend in detail on the well geometry and confining potential. In particular, the prefactor *A*_ex_ *~ w/V*_cavity_, with *w* the polymer effective width and *V*_well_ the well volume. While, in the present case, the polymer is naturally not entirely sterically confined in the well, and can fluctuate into the solution vertically above, this constraint can be relaxed by replacing prefactors appropriate to the steric confinement with prefactors appropriate to the generalized confining potential comprised of steric confinement within the well and field confinement vertically above the well. This field confinement extends above the well and laterally slightly outward due to the curving field funnel. Precisely, the relevant confinement free energy can be computed by solving a mean-field model for a self-excluding chain in the generalized confining potential [45]. To these terms we add a stabilizing free energy −*µξV* (*t*) reflecting the potential energy per unit contour of the chain in the confining electric potential *V* (*t*), with *µ* an electric mobility and *ξ*_*L*_ friction per unit contour. We assume the driving voltage has an offset sinusoidal form *V* (*t*) = *V*_0_ + *V*_*d*_ sin (*ωt*) with *ω* = 2*πf* where *f* is the modulation frequency.

Incorporating all contributions, the full free energy is:

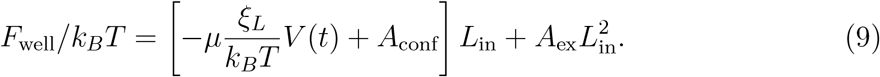

We can then write the equation governing contour exchange:

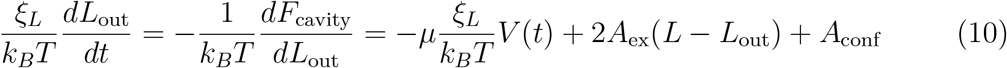

This can be re-parameterized as

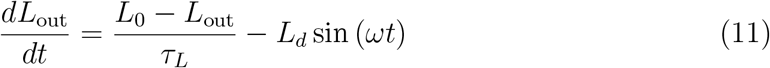

with *τ*_*L*_ ≡ *ξ*_*L*_*/*2*k*_*B*_*TA*_ex_ a relaxation time, *L*_*d*_ ≡ *µV*_*d*_ a driving amplitude, and *L*_0_ the contour present outside the well in the absence of drive modulation (i.e. *V*_*d*_ = 0): *L*_0_ ≡ *L* + (*A*_conf_ − *µV*_0_*ξ*_*L*_)*/*2*A*_ex_.

As contour feeds into the unconfined portion of the chain, the unconfined coil relaxes to the equilibrium values of *R*_*M*_ and *R*_*m*_ consistent with this changing contour. To determine the dependence of *R*_*M*_ and *R*_*m*_ on the unconfined chain contour *L*_out_, we note that *R*_*M*_ and *R*_*m*_ must be proportional to the radius of gyration of the unconfined chain, i.e. *⟨R*_*M*_ *⟩* = *α*_*M*_ *R*_*g*_ and *⟨R*_*m*_*⟩* = *α*_*m*_*R*_*g*_. The radius of gyration can be related to *L*_out_ via the Flory scaling 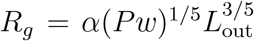 with *P* the chain persistence length and *α* a numerical scaling prefactor. To determine the scaling factors *α*_*M*_ and *α*_*m*_, we cite classic measurements of the principal axes of fluctuating fluorescently stained *λ*− and T4 DNA chains performed by C. Haber *et al* [42], who found that *R*_*M*_ */R*_*m*_ = 2.2, leading to *α*_*M*_ = 0.91 and *α*_*m*_ = 0.414. We model the relaxation dynamics of *R*_*M*_ and *R*_*m*_ using first-order equations with distinct relaxation times *τ*_*M*_ and *τ*_*m*_:

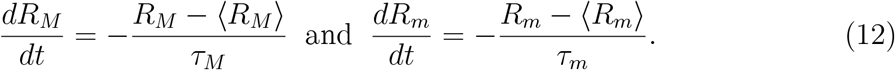

We expect that *τ*_*M*_ *> τ*_*m*_ due to a combination of increased hydrodynamic friction and softer deformation along the major axis.

The system of equations can be simplified by adopting the following dimensionless variables:

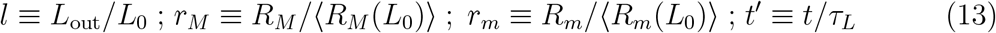

where 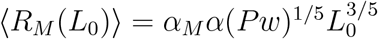 and 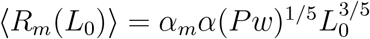. These definitions lead to the reduced system:

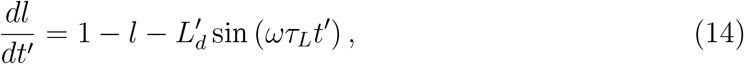

with 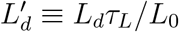 and

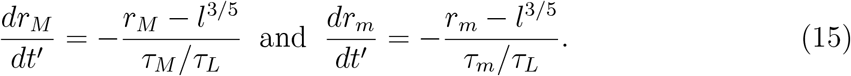

These equations correspond very closely to the action of two first-order filters. The differing relaxation times for *r*_*m*_ and *r*_*M*_ in Eq. 15 lead to different phase offsets when *τ*_*m*_ ≠ *τ*_*M*_. The dynamics of *r*_*M*_ and *r*_*m*_ then involves oscillation at an identical frequency with differing phases, giving rise to closed ellipsoidal trajectories in (*r*_*M*_, *r*_*m*_) space (see Appendix). Figure 8 shows example phase-space loops and behavior of loop geometry for increasing *τ*_*M*_ (Fig. 8A,C) and increasing frequency (Fig. 8B,D). Note that the trajectories collapse to a line for *τ*_*M*_ = *τ*_*m*_ and the behavior is in general non-monotonic (Fig. 8C,D). As *τ*_*M*_ grows relative to *τ*_*m*_ (Fig. 8C), the loop area increases, is maximized and then falls off as slowing dynamics along *r*_*M*_ tilts the ellipse towards *r*_*m*_ and decreases modulation along *r*_*M*_. In the low-frequency limit (Fig. 8D), the loop area is small as *r*_*m*_ and *r*_*M*_ can quasistatically follow the changing equilibrium of the coil outside the cavity. The loop area grows as frequency approaches inverse relaxation times, and then decreases at frequencies surpassing filter cutoffs, with modulation occurring too quickly for the system to follow.

**Figure 8:**
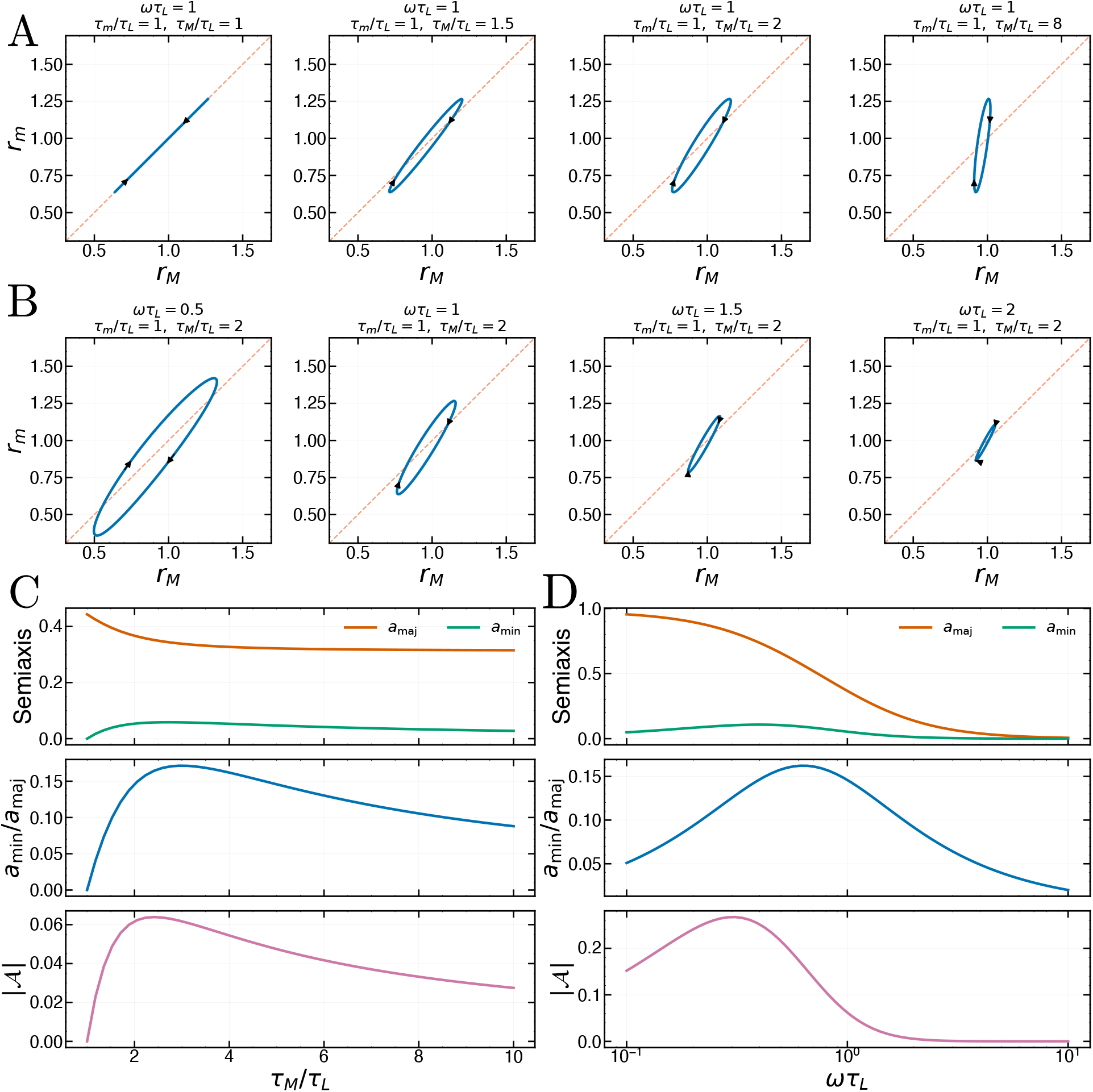
A) Predicted phase-space trajectories for increasing relaxation time along major axis: *τ*_*M*_ */τ*_*L*_ = 1, =1.5, =2 and =8 (left to right) with *ωτ*_*L*_ = 1 and *τ*_*m*_*/τ*_*L*_ = 1. B) Predicted phase-space trajectories for increasing driving frequency: *ωτ*_*L*_ = 0.5, =1, =1.5 and =2 (left to right) with *τ*_*m*_*/τ*_*L*_ = 1 and *τ*_*M*_ */τ*_*L*_ = 1. Note that cycle is traversed in clockwise direction. C) The major and minor axes, major/minor axis ratio and cycle area A as a function of major axis relaxation time with *ωτ*_*L*_ = 1 and *τ*_*m*_*/τ*_*L*_ = 1. D) The major and minor axes, major/minor axis ratio and cycle area A as a function of driving frequency with *τ*_*m*_*/τ*_*L*_ = 1 and *τ*_*M*_ */τ*_*L*_ = 1.

The differing relaxation times for *r*_*m*_ and *r*_*M*_ in Eq. 15 lead to different phase offsets when *τ*_*m*_ ≠ *τ*_*M*_. The dynamics of *r*_*M*_ and *r*_*m*_ then involves oscillation at an identical frequency with differing phases, giving rise to closed ellipsoidal trajectories in (*r*_*M*_, *r*_*m*_) space (see Appendix).

We quantify the two characteristic response timescales by measuring the phase difference between the major and minor conformational modes, represented by 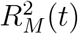 and 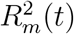, respectively. To determine this phase difference, we extract the complex Fourier coefficients of both time-dependent signals at the applied modulation frequency

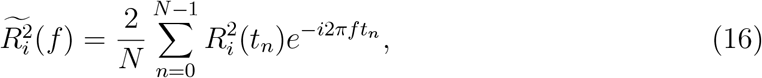

where i=*M, m* and *N* is the experimental number of frames in each experiment. The relative phase difference is then calculated directly from the complex Fourier coefficients as 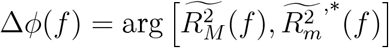 (where the asterisk denotes complex conjugation). This procedure isolates the phase difference associated with the fundamental driving frequency without requiring a fit to the individual time traces. To extract the characteristic response timescales, we fit the frequency dependence of the measured phase difference using our two-mode, first-order relaxation model where the phase difference is given by

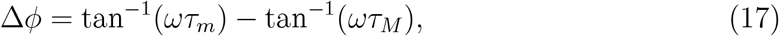

and *τ*_*m*_ and *τ*_*M*_ characterize the responses of the minor and major conformational modes, respectively. The fit yields *τ*_*m*_ = 0.06 ± 0.01 s and *τ*_*M*_ = 0.16 ± 0.04 s (Figure 9A), indicating that the major-axis response is approximately three times slower than the minor-axis response within this model.

**Figure 9:**
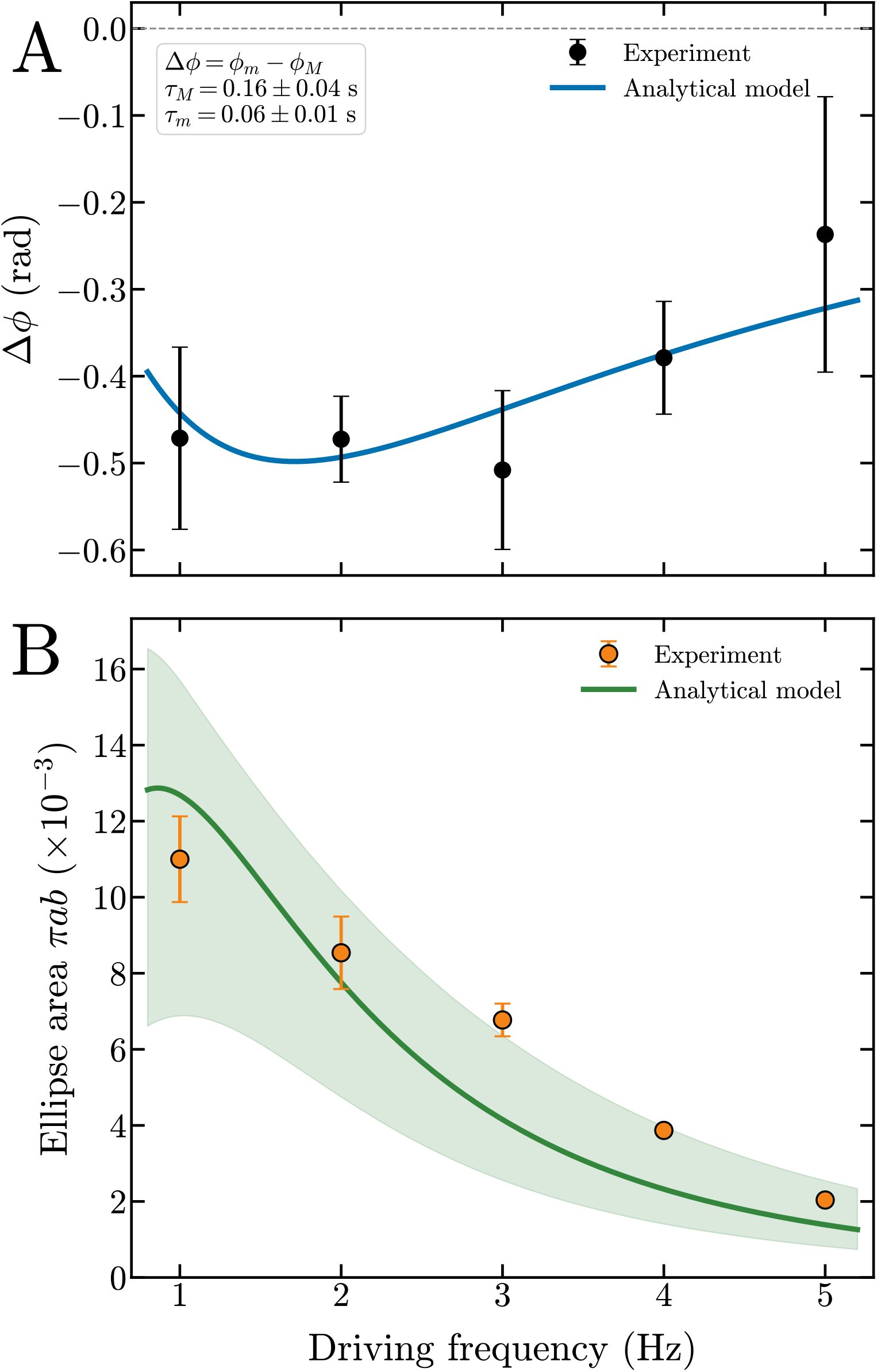
A) Frequency dependence of the phase difference between the major and minor conformational axes, Δ*ϕ* = *ϕ*_*m*_ − *ϕ*_*M*_, fitted using the two-relaxation-time model Δ*ϕ* = tan^*−*1^(2*πfτ*_*m*_) − tan^*−*1^(2*πfτ*_*M*_). The fit yields *τ*_*M*_ = (0.16 ± 0.04) s and *τ*_*m*_ = (0.06±0.01) s. B) Frequency dependence of the conformational loop area, calculated from leastsquares ellipse fits (with fit errors) to the experimental phase-averaged trajectories. The experimentally measured ellipse areas are fitted to the analytical prediction using a single free parameter, the effective amplitude *C*_eff_, while the relaxation times *τ*_*M*_ and *τ*_*m*_ are fixed to the values obtained from the Fourier phase-difference fit.

## 5 Discussion

Analytically, we can obtain an expression for the behavior of the ellipse area in the conformational space as a function of frequency, indcluding the outside-length relaxation time *τ*_*L*_ (Eq. 24). In the limit *ωτ*_*L*_ *≪* 1, the corresponding frequency-dependent factor approaches unity, allowing *τ* to be absorbed into an effective amplitude 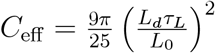 The area then reduces to

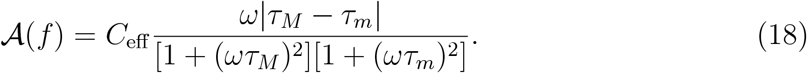

We fit the ellipse area as a function of frequency using this model, obtaining the result shown in 9B, where the lighter green shows the area considering the uncertainties in relaxation of the major and minor axes, respectively. These results suggest that differential relaxation of the conformational axes can be used to qualitatively describe the formation of elliptical trajectories in the conformational phase space. The experimentally measured ellipse areas are also in reasonable agreement with the predicted analytical dependence as a function of frequency in the limit of small relaxation times *τ*_*L*_.

The complete shape of the ellipse as a function of frequency however is not captured by the model. This discrepancy likely arises from the simplifying assumptions of the model, such as describing the response of the major and minor axes through two independent, linear relaxation processes driven by a common periodic excitation. In practice, the conformational dynamics of confined DNA can be more complex, involving nonlinear polymer relaxation, and fluctuations associated with the stochastic motion of the molecule.

Nevertheless, these deviations do not necessarily compromise the Fourier phase analysis that describes the conformational cycling aspect. The phase difference extracted at the fundamental driving frequency characterizes the relative timing of the dominant periodic responses of the major and minor axes, independently of the detailed geometry of the resulting trajectory. Stochastic fluctuations may modify the shape and area of the experimental loops without necessarily producing a corresponding change in the fundamental phase difference. Consequently, while the analytical model is not expected to reproduce every geometrical feature of the experimental trajectories, the phase difference provides a more direct measure of the relative dynamical response of the timescales of the major and minor axes.

The agreement between the measured phase differences and the two-timescale analytical prediction supports the interpretation that there are two distinct effective relaxation times associated with the major and minor axes, in the assumptions of the linear-response model. A more comprehensive description of the cycling dynamics would, however, require explicitly accounting for the coupled motion of the DNA segments inside and outside the nanowell, along with the out-of-plane conformational dynamics along the *z*-direction, which are not directly resolved by our two-dimensional fluorescence measurements. In Equation 15, the contour dynamics depend explicitly on the driving amplitude and the relaxation timescale *τ*_*L*_. Although the applied driving waveform is known, the absence of direct measurements of the DNA contour extending outside the cavity prevents us from independently determining its dynamic response and estimating *τ*_*L*_. These limitations restrict the ability of the analytical model to quantitatively reproduce the complete conformational trajectories.

Furthermore, the experimentally observed DNA conformations are convolved with the microscope’s point-spread function, which affects the apparent size and shape of the molecules. Variations in the axial position of the DNA and fluctuations in the fluorescence signal may introduce additional uncertainties in the measured conformational dimensions, potentially affecting the reconstructed elliptical trajectories, including the apparent tilt and area. Quantitatively reproducing the detailed ellipse shapes would therefore require a more complete description of the three-dimensional conformational dynamics, together with time-resolved measurements of the DNA contour that extends outside the nanowell during each driving cycle. Such measurements are beyond the scope of the present study but could provide additional constraints for future models.

## 6 Conclusion

Here, we have studied the non-equilibrium conformational dynamics of *λ*-DNA molecules under electrokinetic confinement. In the RECON device, molecules are confined electrokinetically by applying a voltage waveform with an offset, which can be toggled by the user in terms of frequency, applied voltage, and waveform. The applied waveform therefore defines an artificial free-energy landscape that can be dynamically perturbed in time. While in passive approaches the DNA molecule acquires a static confined conformation resulting from the interplay between electrostatics and entropy, here the applied external electrokinetic force drives the molecules under confinement to non-equilibrium states. We explored how different electrical waveforms influence the dynamics of the confined molecules. First, we demonstrated how the artificial confinement free energy landscape can be dynamically toggled by varying the frequency, at a constant voltage. We then explore the influence of a stochastic the confinement landscape, and amplitude modulation. We then study in depth the effects of modulation depth and the modulation frequency of the confining potential well on the resulting conformations. The dynamics of the DNA molecules were quantified by tracking the coupled evolution of the major and minor molecular dimensions, *R*_*M*_ and *R*_*m*_, and by visualizing the resulting trajectories in the *R*_*M*_ –*R*_*m*_ conformational phase space. Under amplitude-modulated confinement, we observe hysteresis in the cyclic conformations, and propose a model for such non-equilibrium conformations.

Experimentally, we first kept the voltage fixed, and measured the influence of frequency and confinement geometries on the in-plane conformational extension *R*_*∥*_, from which we extract the major and minor axes of the 2-D projection of the DNA (*R*_*M*_ and *R*_*m*_, respectively). We utilize a series of cavity radii to study the effect of geometry on *⟨R*_*∥*_*⟩/R*_*g*_. Frequencies that are much higher than the diffusion timescale of the molecule (*τ*_*d*_ ≈ 0.18 s) lead to stronger confinement and consequently smaller *⟨R*_*∥*_*⟩/R*_*g*_. Investigating the effect of cavity geometry, when the volume of the cavity is smaller than the *R*_*g*_, the 2-D projection of the DNA is larger, as the molecule occupies a larger volume than the cavity.

Compared to the dynamics of a high frequency, fixed voltage, the stochastic signal leads to higher degrees of molecular excursions from the average, while also leading to slowly decaying relaxation times for the major and minor axes. In the literature, electrokinetic noise has been demonstrated to induce barrier-crossing events and amplify DNA diffusion [43, 44]. While here we demonstrate the influence of the stochastically varying free energy landscape on molecular conformations, the influence of higher and lower noise levels on DNA conformations could be further studied in more detail. For example, one could study the direct influence of effective temperatures in polymer excursions under confinement by modulating the applied noise variance.

We further investigate the confinement using sinusoidally-varying amplitude modulations. First, we find that the degree of amplitude depth can influence the resulting dynamics of the molecule under confinement. While weak modulations (corresponding to 10% − 30% of the maximum voltage) do not result in appreciable time-domain influence over the molecular trajectories in the (*R*_*M*_, *R*_*m*_) phase space, stronger modulations (50% − 120%) lead to hysteresis in the trajectory of the molecular sizes. We propose that such hysteresis in conformational cycling arises from the relaxation timescales associated with the major and minor axes of the polymer coil (i.e. the timescales of *τ*_*M*_ *> τ*_*m*_), leading to a phase offset in the dynamics of the system, and consequently lead to cyclic conformations. Varying the modulation frequency of the applied waveform also leads to hysteresis in the conformational cycles. At low frequencies, the molecules explore a broader region of the (*R*_*m*_*/⟨R*_*m*_*⟩, R*_*M*_ */⟨R*_*M*_ *⟩*) conformational plane, resulting in larger cyclic excursions. As the modulation frequency increases, these excursions become progressively smaller, although the conformational response remains dynamically tunable over the investigated frequency range. This behavior contrasts with the modulation-depth dependence shown in Figure 12. For example, at 50% modulation and 1 Hz, the average molecular conformation remains relatively close to the global mean conformation. In contrast, under 120% modulation, Figure 7 shows that well-defined conformational cycles remain observable even at 5 Hz. Larger modulation depths therefore preserves the cyclic response to higher driving frequencies.

We then developed a simplified semi-quantitative model that couples contour exchange between the nanowell and the external polymer coil to the relaxation dynamics of the molecular conformation. The model treats the DNA contour as partitioned between *L*_in_ and *L*_out_, with the contour length outside the nanowell governed by the competition between confinement, self-exclusion, and the time-dependent electrokinetic trapping potential. This leads to a first-order dynamical equation for *L*_out_, characterized by a contour-exchange relaxation time *τ*_*L*_ and driven by the modulation frequency *f*. Variations in the external contour length determine the instantaneous equilibrium dimensions of the external polymer coil through the Flory scaling relation 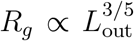. The major and minor conformational dimensions, *R*_*M*_ and *R*_*m*_, are then allowed to relax toward these equilibrium dimensions with distinct relaxation times, *τ*_*M*_ and *τ*_*m*_, respectively. The difference between these timescales introduces a relative phase lag between the two conformational modes, providing a physical mechanism for the emergence of elliptical trajectories in the *R*_*M*_ –*R*_*m*_ conformational phase space. In dimensionless form, the dynamics are governed by the driving parameter *ωτ*_*L*_, the normalized modulation amplitude 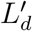, and the ratios *τ*_*M*_ */τ*_*L*_ and *τ*_*m*_*/τ*_*L*_. Experimentally, we determined the two conformational relaxation times by fitting the frequency dependence of the measured phase difference to the analytical two-timescale expression, and subsequently used these timescales to investigate the evolution of the conformational trajectories and enclosed cyclical trajectory areas.

In our theoretical model, *R*_*M*_ and *R*_*m*_ characterize primarily the portion of the polymer coil extending outside the nanowell, whereas the experimentally measured molecular dimensions are obtained from the complete fluorescence image and therefore include both the external coil and the DNA contour remaining inside the underlying well. The current model does not explicitly relate the predicted *R*_*M*_ and *R*_*m*_ to these experimentally measured quantities, which likely contributes to the discrepancies in the ellipse shape. Despite this limitation, the agreement in the overall conformational response supports the physical picture captured by the model. A natural extension of the current model would include incorporating the contribution of the DNA confined within the nanowell and constructing observables from the model that correspond directly to the experimentally measured *R*_*M*_ and *R*_*m*_.

As we demonstrated here, the real time conformation of DNA molecules under confinement in nanocavities can be defined by the user by choosing the applied waveform. In the future, even more complex artificial free energy landscapes can be produced with the RECON device, which could involve different geometries, and waveform characteristics. From a physical point of view, here we considered that the effective electrokinetic funnel that confines the molecules exerts a time-averaged influence over the molecule. Further research is needed to elucidate the full free energy landscape of the electrokinetically confined molecules. Specifically, spatial characteristics of the electric field could locally alter the degree of confinement of the molecules and their conformation. Incorporation of more detailed physics (i.e. electroosmotic flows, dielectrophoresis) could also provide a better description on the organization of the polymers inside electrokinetically-driven nanowells. A more complete modeling of the system could incorporate the distribution of the DNA molecule inside a nanowell considering a mean-field theory, specifically under the influence of an external electric field [46]. A better calibration of the effective electric field inside a nanowell would require further continuous measurements of the current through a single nanocavity. This would enhance understanding of the time-dependent electrokinetic confinement devices.

## 7 Appendix

Here we briefly sketch the details of the solution of the system of equations 14 and 15. Eq. 14 is a linear first order system and leads, after any transient has decayed away, to:

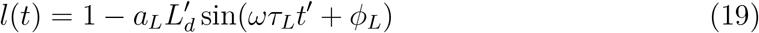

with 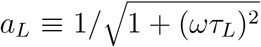 a low-pass filter acting on the amplitude and the phase offset *ϕ*_*L*_ ≡ − tan^*−*1^(*ωτ*_*L*_). If we assume the driving amplitudes are sufficiently small so we can linearize 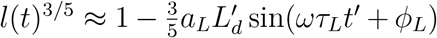, then the solutions for the principal axes are simply fed through another linear first order system leading to:

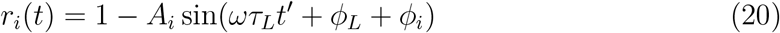

with 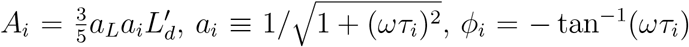 and *i* = *m, M*. As the solutions for *r*_*m*_ and *r*_*M*_ have the same driving frequency but different phases, the trajectory is an ellipse. Eliminating 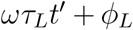 between solutions for *r*_*m*_ and *r*_*M*_ leads to the closed phase-space trajectory:

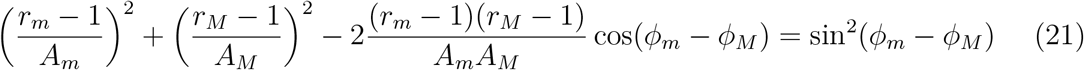

This collapses to a line in the case *ϕ*_*m*_ = *ϕ*_*M*_.

A more transparent and systematic way to view the elliptical phase-space geometry is to consider it arising from a linear transformation of a circle via a combination of rotation and stretching. Let 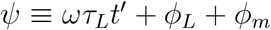 and *δ* ≡ *ϕ*_*M*_ − *ϕ*_*m*_. Then define *y* ≡ 1 − *r*_*m*_ = *A*_*m*_ sin(*ψ*) and *x* ≡ 1 − *r*_*M*_ = *A*_*M*_ sin(*ψ* + *δ*) = *A*_*M*_ [cos *ψ* sin *δ* + cos *δ* sin *ψ*]. The action of the conformational dynamics is then to map the circular trajectory (sin *ψ*, cos *ψ*) to an elliptical trajectory (*x, y*) via the transformation matrix:

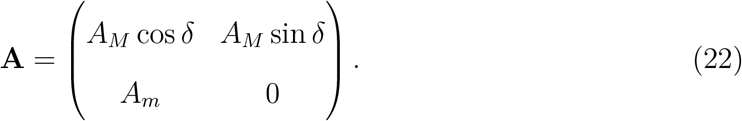

All properties of the elliptical trajectory can be obtained from **A**, including the semiminor (*a*_min_) and major (*a*_maj_) axis which are related to the square root of the eigenvalues of the matrix **S** = **AA**^*T*^. The ellipse area quantifying hysteresis *A* = *πa*_min_*a*_maj_ = *A*_*m*_*A*_*M*_ | sin(*δ*)|. For reference, we give the explicit forms for *a*_maj_ and *a*_min_:

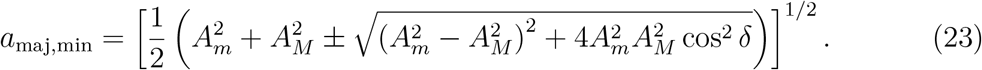

The equation for the ellipse area has the explicit form in terms of frequency:

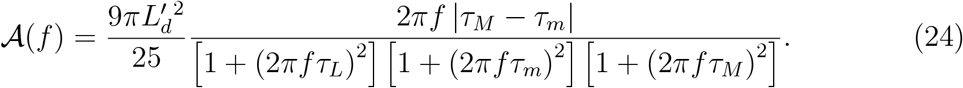

At very low frequency *A* is minimal, as the major and minor axes have sufficient time to adjust to the changing out of cavity contour. The hysteresis grows as the relaxation times are approached, then decreases at high frequency as the system can no longer follow the rapid dynamics (low-pass filtering).

## 8 Supplementary material

### 8.1 Conformational distributions under constant voltage confinement

In Figure 10, we present the distributions of *R*_*M*_ and *R*_*m*_ for fixed voltage (*V* = 5 V), for varying frequencies between *f* = 1Hz− 1MHz. We investigate the influence of varying the cavity sizes (*R* = 250 nm in blue, *R* = 300 nm in yellow, *R* = 350 nm in green, *R* = 400 nm in red) in the average *R*_*M*_ and *R*_*m*_. Following the discussion presented in the main text, frequencies that are above the relaxation timescales of the DNA in the nanowells (*f >* 10 Hz) lead to decreasing averages of *R*_*M*_ and *R*_*m*_, while also bringing averages closer together. Notably, increasing the cavity radius also results in tighter distributions and smaller differences between the two axes. As the confining volume increases, a larger fraction of the DNA contour can be accommodated within the nanowell, reducing the need for the molecule to extend beyond the cavity and resulting in a more isotropic conformation.

**Figure 10:**
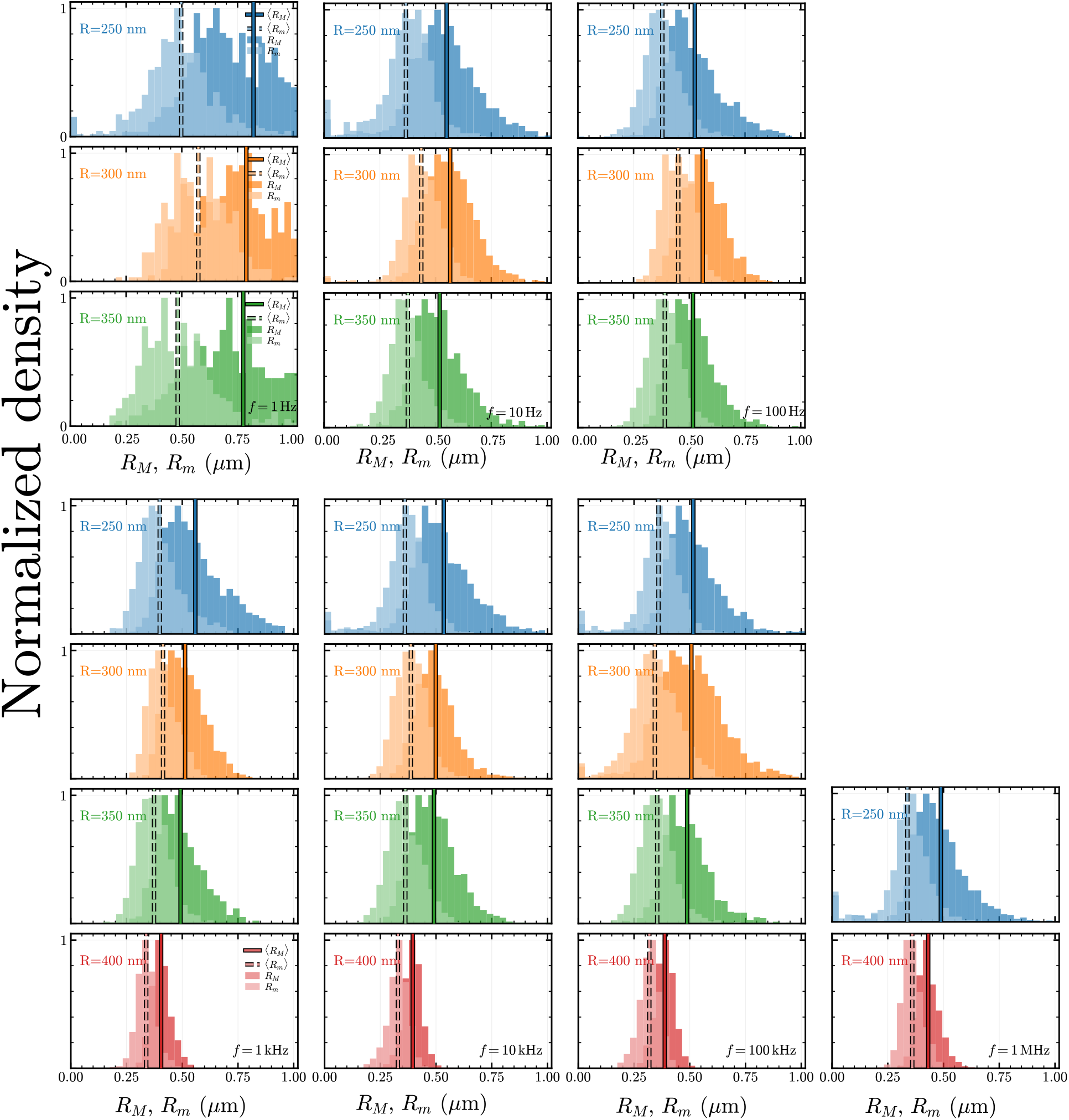
A) Histograms showing the normalized density of the major and minor axes of the radius of gyration tensor, *R*_*M*_ and *R*_*m*_, for different cavity radii (*R* = 250 nm in blue, *R* = 300 nm in yellow, *R* = 350 nm in green, and *R* = 400 nm in red) and different frequencies, at fixed voltage voltage (*V*_min_ = 0 V, *V*_max_ = 5.0 V). The dashed and outlined vertical lines giving the averaged radii.

### 8.2 Relaxation timescalesfor different geometries

In Figure 11, we present the results for the autocorrelation of the *R*_*∥*_ for various cavity diameters. We note that the relaxation timescales are very short for all cavity diameters, and do not present any time-domain features regarding *R*_*∥*_.

**Figure 11:**
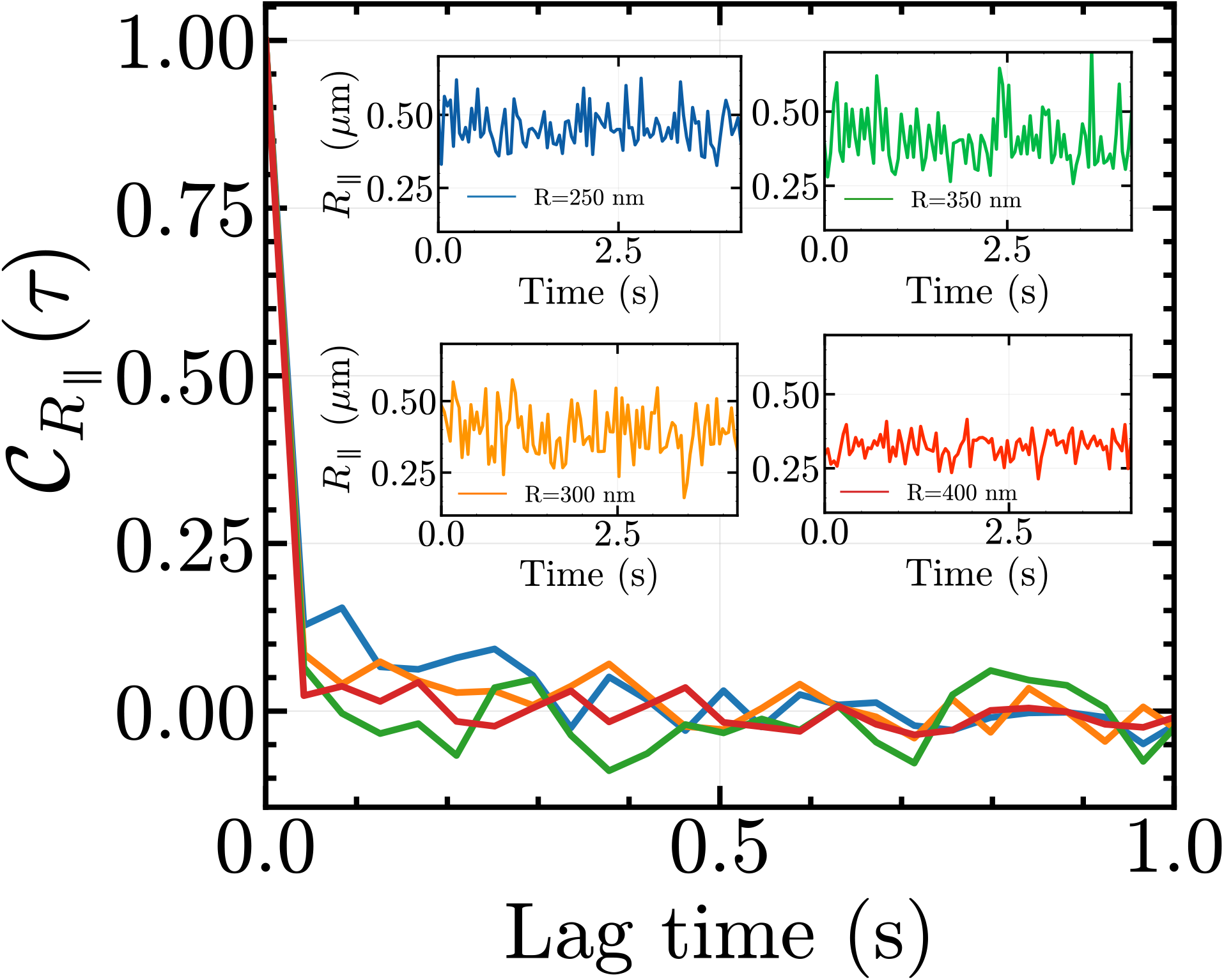
Autocorrelation of *R*_*∥*_ for different cavity radius, at constant voltage and constant frequency *f* = 100 kHz. There is no time-signature, showing the modulation of *R*_*∥*_ at high frequencies. At lower frequencies (i.e. 1–10Hz), we do see modulation.

### 8.3 Conformational cycling as a function of AM percentage

In Figure 12, we show the amplitude-dependent conformational response of confined DNA under amplitude-modulated electrokinetic confinement for modulation depths of 10%, 20%, 30%, 50%, 100%, 110%, and 120%, with modulation frequency *f* = 1 Hz. complementary to the amplitudes presented in the main text. We observe that the degree of conformational excursions increases as the modulation depth increases. From 50% amplitude depth, we observe cyclic conformational trajectories in the (*R*_*M*_, *R*_*m*_) phase space. This dynamics is also reflected by the sinusoidal behavior of *R*_*m*_, *R*_*M*_ and the normalized in-plane molecular extension *R*_*∥*_. We present in Figure 13 the autocorrelation function 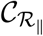 for the various amplitude modulation depths, fitted with an exponential decay (10%, 20%, and 30%), and a exponentially decaying sinusoidal model (50%, 100%, 110%, and 120%).

**Figure 12:**
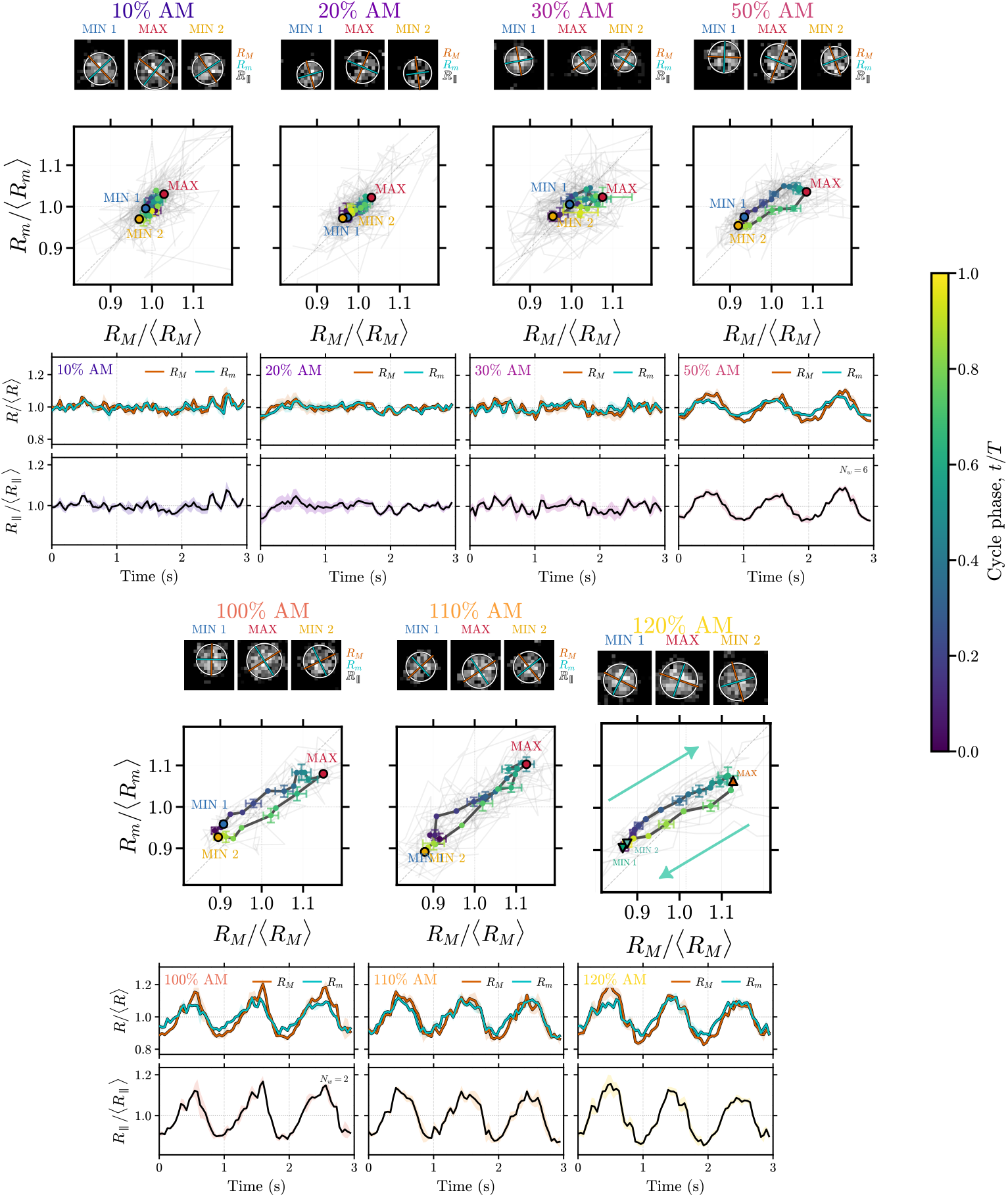
Amplitude-dependent conformational response of confined DNA under amplitude-modulated electrokinetic confinement. A) Applied voltage waveforms for modulation depths of 10%, 20%, 30%, 50%, 100%, 110%, and 120%. The upper and lower envelopes indicate the instantaneous modulation limits, *V*_max_ and *V*_min_, over three modulation periods, *t/T*. B) Corresponding conformational trajectories in the normalized *R*_*M*_ –*R*_*m*_ phase space for *n* = 10 cycles along with the averaged trajectory (pale gray indicates underlying single-cycle trajectories; black bold curve indicates average). Representative fluorescence images at the starting point where signal is at its maximum and coil size in minimum (MIN 1), signal is minimized and coil size is maximum (MAX), and return to state of minimum coil size (MIN 2) are shown above each trajectory. Colored points along the averaged trajectory indicate progression through the modulation cycle according to the normalized cycle phase *t/T*, with dark blue indicating the start of the cycle, red the middle maximum point on the cycle and bright yellow indicating the cycle end. Error bars correspond to standard error of the mean of each point within the cycle and indicate the variability of the ensemble-averaged conformational coordinates at these positions. Increasing modulation depth produces progressively larger excursions in *R*_*M*_ */⟨R*_*M*_ *⟩* and *R*_*m*_*/⟨R*_*m*_*⟩*, with the forward and backward portions of the cycle defining a closed conformational trajectory.

**Figure 13:**
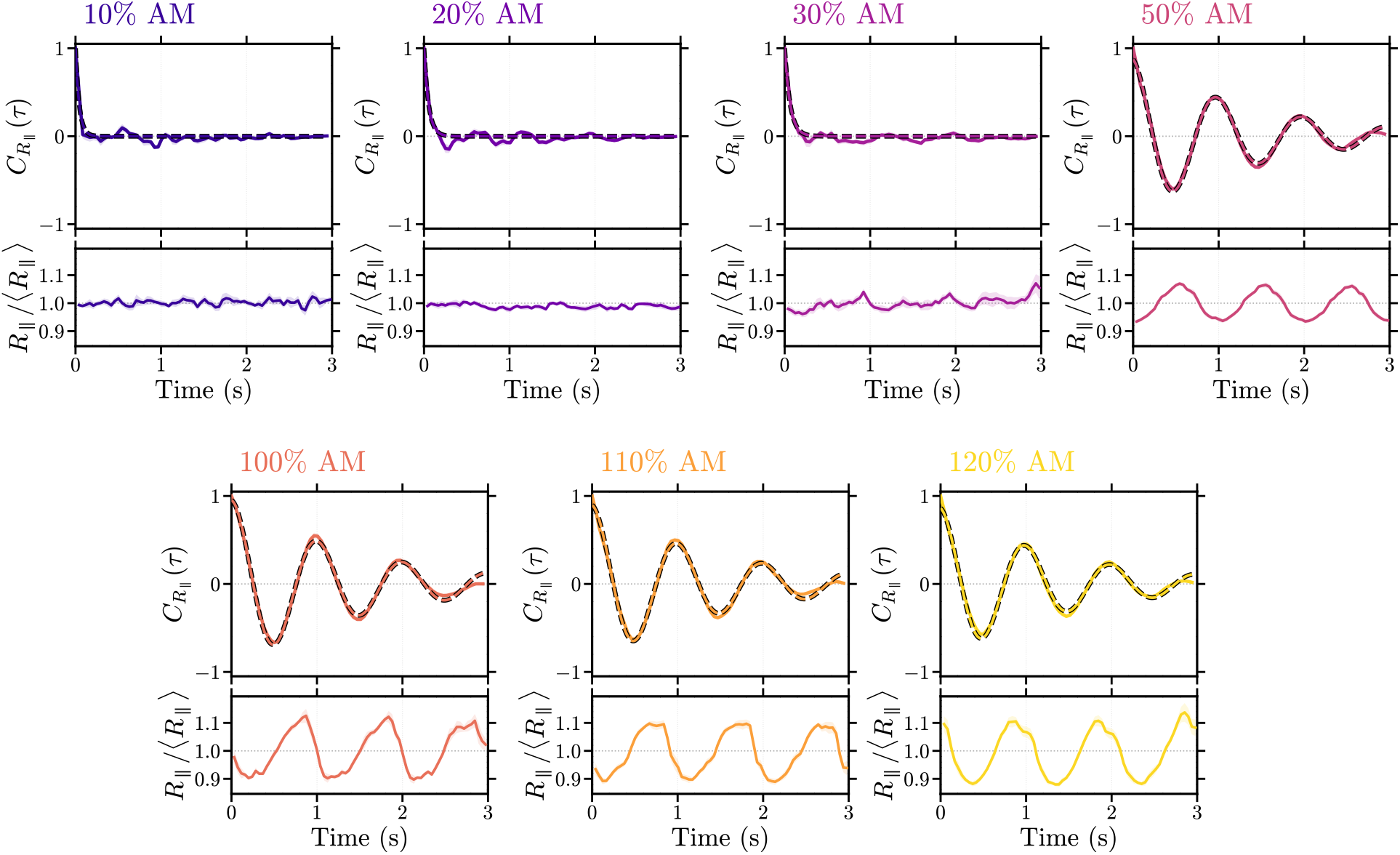
Autocorrelation functions of the normalized in-plane molecular extension, *R*_*∥*_*/⟨R*_*∥*_*⟩*, as a function of amplitude-modulation depth. Colored curves and shaded regions indicate the ensemble-averaged 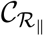 and SEM, respectively, while dashed curves show the corresponding fits. The average *R*_*∥*_ response over the same 3 s interval is shown below each 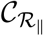. Increasing modulation depth produces a transition from rapidly decaying correlations to a pronounced oscillatory response synchronized with the 1 Hz modulation.

